# Mosquito sound communication: are male swarms loud enough to attract females?

**DOI:** 10.1101/2020.09.01.277202

**Authors:** Lionel Feugère, Gabriella Gibson, Nicholas C. Manoukis, Olivier Roux

**Affiliations:** MIVEGEC, Univ. Montpellier, IRD, CNRS, Montpellier, France; Natural Resources Institute, University of Greenwich, Chatham, Kent ME4 4TB, UK; Tropical Crop and Commodity Protection Research Unit, Daniel K. Inouye US Pacific Basin Agricultural Research Center, US Department of Agriculture- Agricultural Research Service, 64 Nowelo St. Hilo Hawai‘i USA 96720; Institut de Recherche en Sciences de la Santé (IRSS), 01 BP 545 Bobo-Dioulasso 01, Burkina Faso

**Keywords:** *Anopheles gambiae*, free-flying mosquitoes, long-range hearing, mating swarm, mosquito hearing, speciation

## Abstract

Given the unsurpassed sound-sensitivity of mosquitoes among arthropods and the sound-source power required for long-range hearing, we investigated the distance over which female mosquitoes detect species-specific cues in the sound of station-keeping mating swarms. A common misunderstanding, that mosquitoes cannot hear at long-range because their hearing organs are ‘particle-velocity’ receptors, has clouded the fact that particle-velocity is an intrinsic component of sound whatever the distance to the sound source. We exposed free-flying *Anopheles coluzzii* females to pre-recorded sounds of male *An. coluzzii* and *An. gambiae s.s.* swarms over a range of natural sound-levels. Sound-levels tested were related to equivalent distances between the female and the swarm for a given number of males, enabling us to infer distances over which females might hear large male-swarms. We show that females do not respond to swarm sound up to 48 dB SPL and that louder SPLs are not ecologically relevant for a swarm. Considering that swarms are the only mosquito sound-source that would be loud enough to be heard at long-range, we conclude that inter-mosquito acoustic communication is restricted to close-range pair interactions. We also showed that the sensitivity to sound in free-flying males is much enhanced compared to that of tethered ones.

## Introduction

One-on-one male-female auditory interactions in mosquitoes have been shown to be related to pre-mating behaviour in at least four species of medical importance (*Anopheles gambiae s.l.*, *Anopheles albimanus*, *Aedes aegypti* and *Culex quinquefasciatus*), plus *Toxorhynchites brevipalpis* and *Culex pipiens (1–10)*, as well as in other dipteran flies *(11)*. It is assumed that the hearing distance between a male and a female is limited to a range of a few centimetres to ~ 10 cm *(12, 13)*. However, although their auditory organs are optimized for close-range hearing, they are not restricted to a given hearing distance *(14)*, because they are sensitive to an intrinsic component of sound *(15, 16)*. Consequently, males have been shown to respond to artificially-loud sound levels of played-back single female flight-tones metres away from the sound source *(16)*. Thus, the debate about hearing distance should be strictly linked to sound-source power and the biological relevance of the sound-source in the field. In other words, is long-range inter-mosquito sound communication *(16)* only possible in the laboratory, or does it also occur under natural environmental conditions? From existing results, it is reasonable to assume that to be heard at distances greater than ~10 cm the source of mosquito sound must be more powerful than that of an individual mosquito. Species of mosquito that form mating swarms can produce a relatively loud sound, easily discernible to the human ear a few metres away *(17)*, by forming relatively dense station-keeping aggregations *(18)*, consisting of up to thousands of males *(19–21)*. This raises the hypothesis that a female can be attracted from a distance to swarm sounds produced by males in established swarms.

Electrophysiology measurements show that mosquito auditory organs are the most sensitive among arthropods when exposed to the sound of an opposite-sex individual *(13)*, with females generally slightly less sensitive than males *(1, 16*; but see *22)*. Behaviour studies demonstrate that, although females have not been shown to move toward the sound source of an individual male (phonotaxis), females of at least one mosquito species uses phonotaxis to locate a blood feeding host *(23)* and females of several mosquito species alter their wingbeat frequency when exposed to male sound *(1, 3, 24)* probably to hear the male better *(3, 6)*. An important lacuna in the literature remains: can a single female hear the sound of an entire swarm of conspecific males?

The two species of the *An. gambiae s.l*. complex we worked with (*An. coluzzii* and *An. gambiae s.s.)* are African malaria vectors and are under-going speciation *(25)*. These species are found in sympatry and mainly mate assortatively. Subtle differences in swarming behaviour between these closely related mosquitoes can minimize hybridization. Female auditory detection of a con-specific swarm of males at long range could increase the female’s likelihood of locating and being inseminated by a male of the same species. A female might recognize a species-specific sound signature at long-range before males of any other species could hear, chase and mate with her. Species-specific acoustic cues in *An. coluzzii* and *An. gambiae s.s*. have been reported based on studies of single male or male-female pair interactions. Laboratory-based research characterizing the flight tones of single males flying “randomly” in cages found no significant differences between the fundamental frequencies of *An. coluzzii* and *An. gambiae s.s*., although significant differences were found in the second harmonic amplitude *(26)*. In a separate study, the rapid wingbeat frequency modulations associated with mating *(6, 8, 9)* were found to be similar when males of both species were exposed to pure tones mimicking the female’s fundamental wingbeat frequency *(27)*. However, another study of the patterns of flight tone interactions between a tethered male and a tethered female of closely related species of *An. gambiae s.l*. found that frequency-matching occurred more consistently within pairs of the same species than in hetero-specific pairs *(4)*, and frequency matching was shown to be associated with mating success in *Aedes (8)*. These close-range studies are interesting, but they beg the question as to what occurs in the lead-up to close-range interactions. To our knowledge, the response of females to the species-specific sound of distant male swarms has not been tested quantitatively yet.

Accordingly, our hypothesis is that uninseminated *An. coluzzii* female mosquitoes detect distant sounds of swarming con-specific males at natural sound levels and respond to species-specific cues in the swarm sound. In Burkina Faso, we recorded ambient sound in the field near naturally swarming *An. coluzzii* males to determine whether any other animal or environmental sounds were present that could hide/mask swarm sounds: mosquito sounds stand out against ambient noise at least 3 m from the swarm (see Figure S1 and supplemental information section ‘Mosquito sounds stand out against ambient noise at least 3 m from the swarm**’**). Thus, we decided to test our hypothesis under laboratory conditions using sound levels derived from 1) calibrated sound recordings of swarms and 2) model of swarms and their acoustics in order to rigorously extrapolate behavioural results to larger swarms and various distances that would have been difficult to achieve with real swarms in the laboratory. This application of acoustics theory, including an accurate reproduction of the particle-velocity field and the estimates of acoustic measurement uncertainties, served to validate our conclusions.

## Materials and Methods

### Experimental principle based on behaviour assay and acoustic propagation theory

We conducted behavioural experiments in an environmentally controlled laboratory fitted with a soundproof chamber (Figure 1), by presenting sound recordings of swarming males to free-flying females (see Supplementary Methods section ‘Generation of sound stimuli’ and ‘Sound pressure level’). Free-flying uninseminated females were released in a swarming arena (L x W x H = 1.8 m x 1.7 m x 2 m) that provided the visual cues (see Supplementary Methods section ‘Environmental conditions in soundproof chamber’) to initiate swarming flight (figure-of-eight loops) over a visual marker, effectively confining them to a volume of 0.06 m^3^ and within a fixed distance from the source of male swarm sound (Figure 2 A). Instead of changing the distance between the test female and the male swarm, we used a range of sound levels to mimic a range of distances between a female and swarming males, altering the apparent distance *r*_*i*_ between the female and the sound-source ‘image’ of the played-back swarm according to acoustic propagation theory (see Supplementary Methods section ‘Formulae relating sound level and distance’):

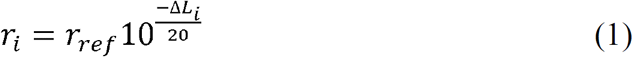

with: *r*_*ref*_ = 0.9 m, distance between the speaker and the swarm centre; Δ*L*_*i*_ being the SPL difference between *r*_*i*_ and *r*_*ref*_. for a 70-male swarm.

**Figure 1.**
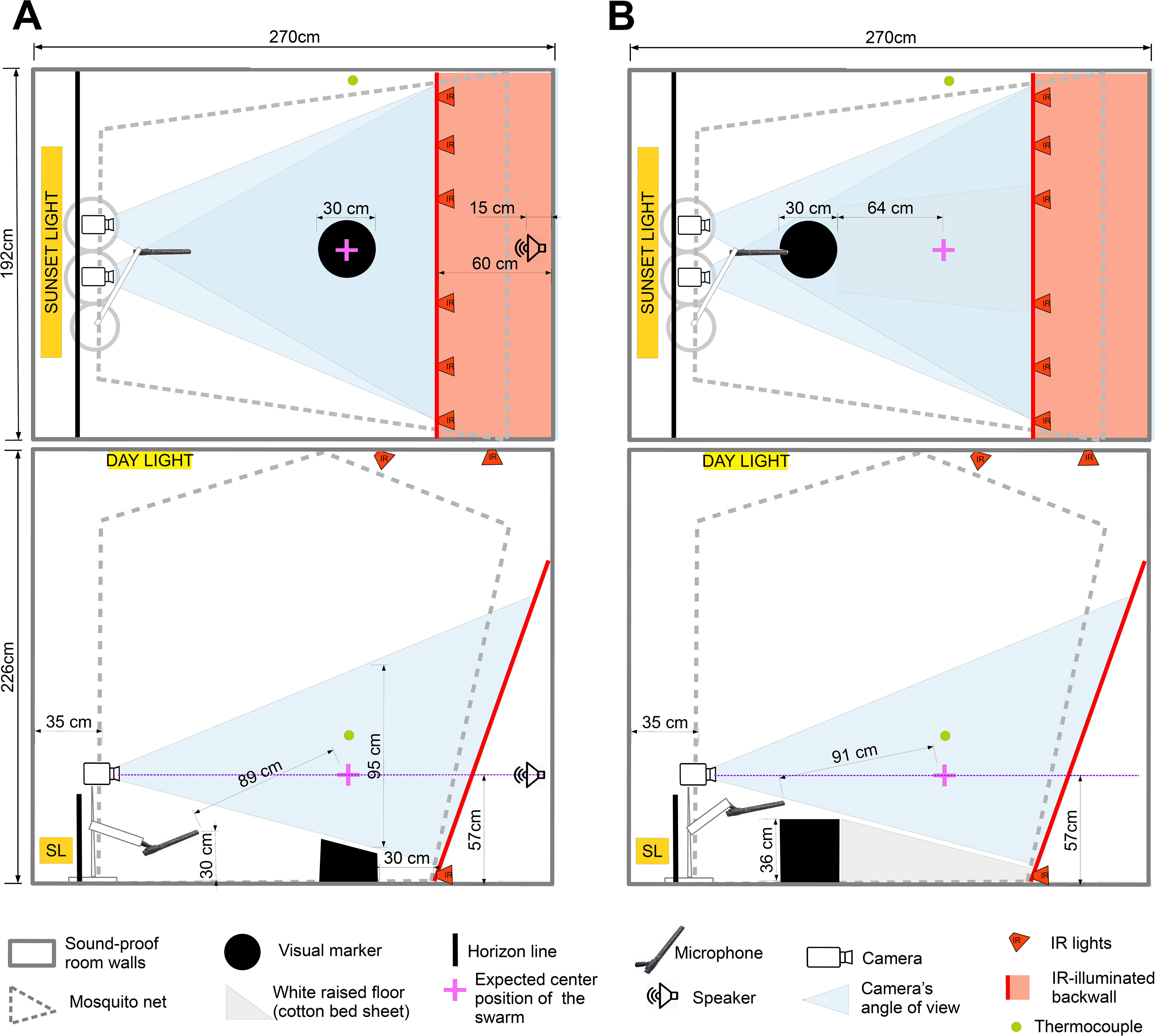
Soundproof chamber setup for recording sound and video of mosquito behaviour. Bird’s-eye and side views of soundproof chamber. Two IR-sensitive cameras fitted with IR pass filters tracked flying mosquitoes as black silhouettes against evenly lit IR-background. Separate lighting system provided gradual semi-natural dusk visible to mosquitoes, consisting of dispersed dim white lights on ceiling and ‘sunset’ lighting below horizon (opaque wall ~40 cm tall). A microphone recorded flight sounds of mosquitoes swarming directly above black swarm marker. A thermocouple (85 cm above ground level) recorded temperature at ~ mean swarm height. Differences between setups for the two species was necessary to accommodate species-specific differences in positioning of swarming flight in relation to swarm marker *(29)*. **(A)** Setup to record sound and flight of *Anopheles coluzzii*, for sound stimulus recording and behavioural experiment. A speaker located behind IR-illuminated thin-cotton sheet, outside net enclosure played back sound stimuli. **(B)** Setup to record sound of *Anopheles gambiae s.s*., for sound stimulus recording only.

**Figure 2.**
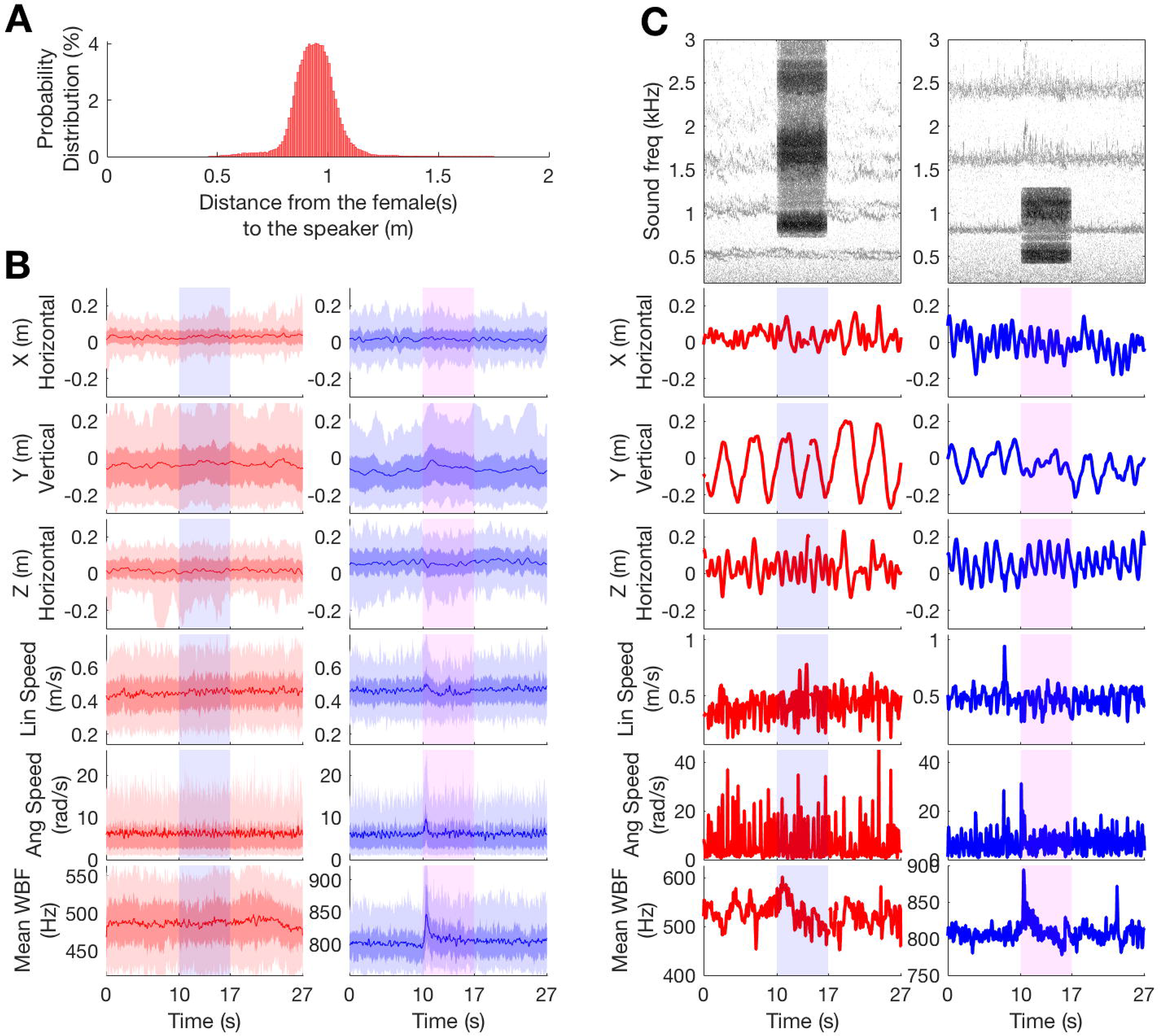
Flight and sound responses of females and males to sound stimuli. Female (red) and male (blue) flight-characteristics and wingbeat-frequencies before, during and after playback of male (blue rectangle) or female (red rectangle) sound stimuli. **(A)** Probability distribution of distance between a female and the speaker during sound stimulus playback; 95% of distances were between 72 cm and 113 cm, with a mean and median of 94 cm. This distance interval was used to estimate the uncertainties of the acoustic prediction in Table S1 and Figure 5. **(B)** *Anopheles coluzzii* response to highest sound-level *An. coluzzii* and *An. gambiae* sound-stimulus over 27 s of recording. Stimulus was played-back 10 s from beginning of flight recording and lasted 7 s (red or blue rectangular shading). First five rows show flight parameters (relative ‘XYZ’ position, plus linear and angular flight speeds). ‘Z’ dimension represents relative distance to the speaker (located 0.9 m from *Z*=0). Last row shows mean wingbeat frequency (WBF) of 1^st^ harmonic. Darkest coloured lines represent running median, darkest areas represent second and third quartiles and light areas represent the 90^th^ percentile of data. Distribution of flight coordinates and velocities were recorded for 149 female tracks and 104 male tracks, and the WBF distribution plot is based on mean WBFs over the number of mosquitoes per fly group (100 female-groups and 61 male-groups). No clear apparent response was observed in females, whereas for males, linear and angular speed and wingbeat frequency clearly increased in response to the sound stimulus onset, plus a slight tendency to increase the flight height was evident. **(C)** Same as B (with the exception of the spectrogram), but with a single example per plot. First row shows spectrograms of sound recordings before, during and after the sound stimulus. The color gradient represents the sound level given a frequency and a time (the darker the color, the louder the frequency). Spectrogram in the first column displays a live *An. coluzzii* female exposed to *An. coluzzii* male sound between 10th and 17th s (Video S1), while the spectrogram in the second column displays a live *An. coluzzii* male exposed to the two first-harmonics of the *An. gambiae* female sound (Video S2). Periodic flight pattern, typical of swarming behaviour, is evident for males and females in ‘XYZ’ plots. See Figure S3 for a superimposition of all tracks from the 48 dB *An. coluzzii* stimulus.

Finally, the measured results were extrapolated to estimate how far away (*r*_*i,N*x70_) a female mosquito can hear a swarm with *N* times more males (see Method section ‘Acoustic assumptions to model a swarm at long-range’ below and Supplementary Methods section ‘Formula relating hearing distance and number of individuals in the swarm’):

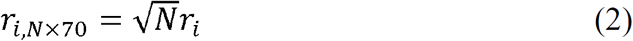

Values are presented in Table S1 for a 300, 1,500, 6,000 and 10,000-male swarm and Figure 3 summarizes the experimental principle and the raw results.

**Figure 3.**
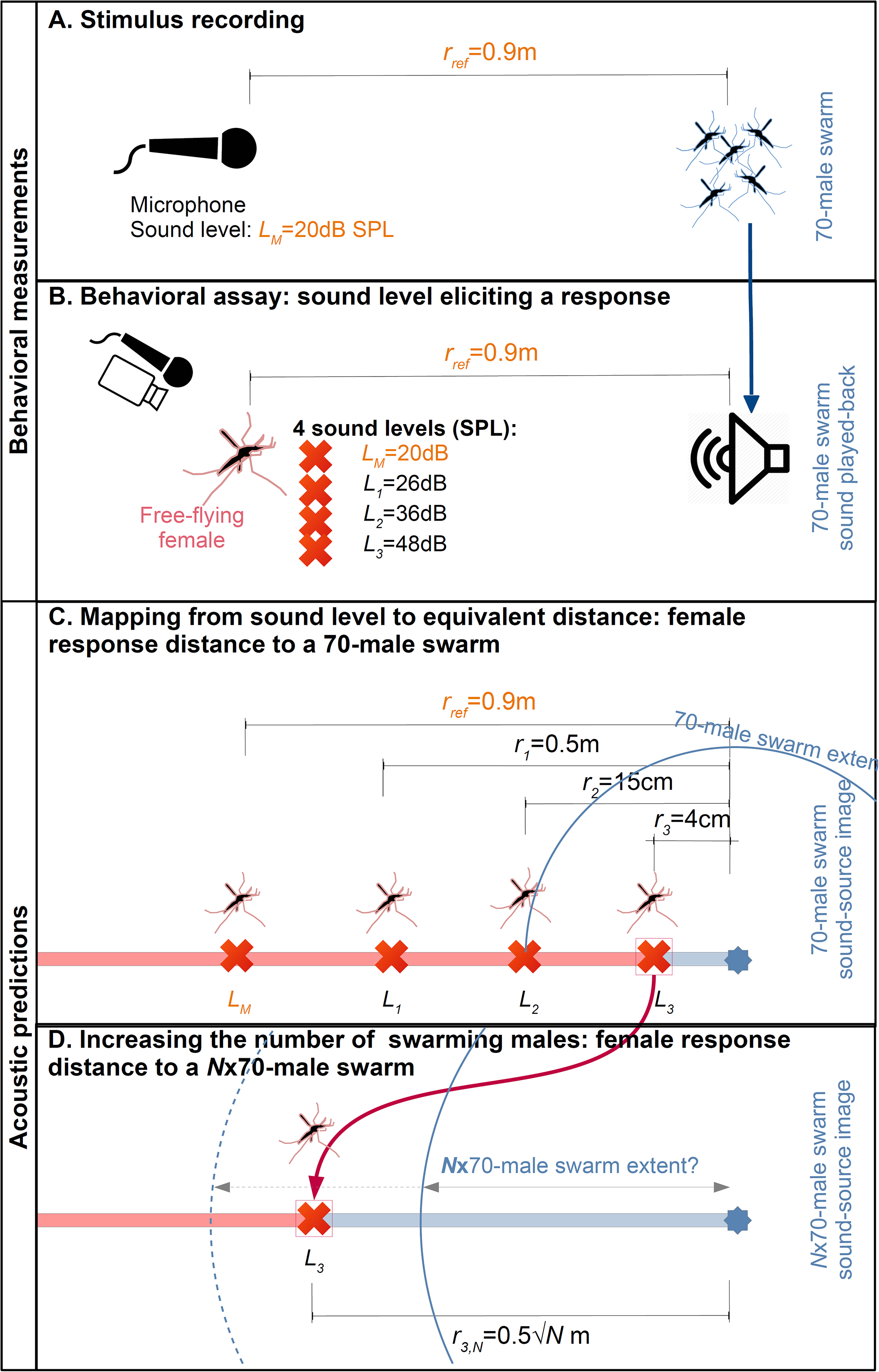
Steps to evaluate the distance a female mosquito can detect the sound of an *An. coluzzii* male swarm of a given number of individuals. This schematic explanation shows how methodologies from behavioural assays (‘measurements’) and acoustic theory (‘predictions’) were employed in this study, based on details for *An. coluzzii* sound stimuli. The same procedure was repeated with sound stimuli of *An. gambiae s.s*. and the reciprocal experiment was performed with males exposed to sound stimuli of a female-swarm for both species. **(A)** First, the reference stimulus (sound of 70 males swarming) was recorded at 0.9 m from the male swarm, producing a sound pressure level of 20 dB SPL. **(B)** Second, this stimulus was played-back to 1-5 swarming (station-keeping) females in free-flight at four different sound levels (20, 25, 36 and 48 dB SPL) as measured at the mean females’ distance to the speaker (see Figure 2 A). None of them triggered a response in females. **(C)** Third, assuming the swarm sound emitted from the speaker to be a point source, and given the natural sound level of a 70-male swarm (*L*_*M*_) at a distance of 0.9 m (*r*_*ref*_), we can compute the distance to a similar swarm corresponding to the other three sound levels (see Supplementary Methods) and compare it to the swarm radius. **(D)** Fourth, the effect of multiplying the number of swarming males per N over the loudest stimulus is predicted (see Supplementary Methods).

### Control of distance between live mosquito and playback speaker

To establish fixed distances between the sound source and free-flying females, we exploited female swarming behaviour; in the absence of male mosquitoes, uninseminated females swarm over a floor marker in flight patterns similar to those of conspecific males *(28, 29)*. Accordingly, we constructed a flight arena that provided visual cues that stimulated females to fly in elliptical loops over a stationary swarm marker, effectively confining them within a limited area of the flight arena *(28, 29)*, which enabled us to assess whether or not a female responded to the sound stimulus of the playback of swarming males at a controlled sound level. The speaker (Genelec 8010A) that reproduced the males’ swarming flight tones was placed 0.9 m from the centre of the swarm marker. A few females (< 15) at a time were released in the flight arena, and periodically 1 to 5 females were stimulated by the visual characteristics of the marker to switch from ‘random’ flight to swarming flight. Their flight positions were recorded by 3D-tracking Trackit Software *(30)* (Figure 2 B, Figure 2 C) which enabled us to determine the distance between a mosquito and the speaker emitting mosquito sound (0.9±0.2 m, 95%-CI, Figure 2 A).

### Physical sound quantities produced by a speaker and sensed by mosquitoes

Like any sound-source, a speaker creates both a pressure field and a particle-velocity field. We monitored the sound level of our swarm sound playbacks by recording the sound pressure level (SPL), while mosquito hearing organs are sensitive to particle velocity levels (SVL) *(31)*. These two quantities are equal only far from the sound source. Considering the frequency content of male swarm sound (no frequency components below *f* = 745 Hz), we can calculate that for *r >* 15 cm, SPL and SVL are equal with an error less than 1 dB (see Supplemental Methods section ‘Relationship between particle-velocity and pressure levels’). Then in our case, monitoring the swarm playback SPL is equivalent to monitoring SVL.

### Acoustic assumptions to model a swarm at long-range

The density of a swarm is far greater in the centre than at the periphery *(18)* (Figure S2). Therefore, for the purposes of this analysis, we considered the swarm to be a point source that radiates spherically in all directions (neglecting the sound reflection on the ground or any nearby object). This approximation can be used if the swarm radius remains relatively small compared to the distance between the female and the swarm centre. Swarms can be ovoid *(29, 18)*, but this is not an issue for our point-source assumption, because the oval dimension is often perpendicular to the female-to-swarm spatial axis, so each swarming male equally contributed to the radiated swarm sound toward the female at long range. Our recorded swarm is composed of 70 males and all the other acoustic predictions are performed with swarms of hundreds to thousands of individuals, which would considerably attenuate any effect of individuals and then forming a single sound-object entity. In addition, we will see in the discussion that our hypothesis of considering a monopole source rather than multiple dipole sources is conservative for our results.

### Experimental design

For each replicate (one per day, August-September 2018), about fifteen 3-6 days-old uninseminated females were released the day prior to experiments at ~ 18h00 in the sound recording flight arena and left to fly freely until the end of the experiment. At 15h00, after the ceiling lights had dimmed to the lowest intensity, the horizon light completed a 10 min dimming period and then was kept at a constant dim light intensity until the experiment was finished. When at least one female started to swarm robustly over the marker, a first sequence of sound stimuli was played. Each of the subsequent sequences were played immediately following the last if the previous female(s) was still swarming or as soon as at least one female started swarming. The experiment was ended when the maximum number of stimuli sequences (10) was reached or after 50 min of constant horizon light. Females were then collected and removed from the flight arena. A new group of ~15 mosquitoes were released in the soundproof chamber, to be used for a new replicate the next day.

### Subject details

All experiments were performed with two sibling species in the *Anopheles gambiae s.l*. Giles species complex: *An. gambiae s.s*. Giles and *An. coluzzii* Coetzee & Wilkerson. Colonies of the two species were established at the Natural Resources Institute (NRI), University of Greenwich (UK) from eggs provided by the Institut de Recherche en Sciences de la Santé (IRSS), Burkina Faso. *Anopheles coluzzii* eggs were obtained from a colony established in 2017 from wild gravid females collected from inhabited human dwellings in Bama, Burkina Faso (11°23’14″N, 4°24’42″W). *Anopheles gambiae s.s*. eggs were obtained from a colony established at IRSS in 2008 and renewed with wild material in 2015 from Soumousso, Burkina Faso (11°00’46”N, 4°02’45”W). Females were identified to species level by PCR *(32)*. The NRI colonies were kept in environmentally controlled laboratory rooms with a 12h:12h light:dark cycle (lights went off at 15h00), >60% relative humidity and ~24-26°C. Larvae were fed Tetramin® fish-flakes and rice powder. Adult males and females were separated < 12h post-emergence to ensure females were not inseminated, and fed a solution of 10% sucrose in an isotonic saline *ad libitum*.

### Statistics

Flight trajectories were measured by the 3D-tracking software *(30)* and wingbeat frequencies were extracted from the sound recording using Matlab (see Supplemental Methods section ‘Response parameters’, Figure 2 B, Figure 2 C, Figure S3). We were not able to discriminate between mosquitoes from their wingbeat frequencies when swarming in a group, so for each sound parameter, values were computed for the whole tested group of 1-5 females or of 1-6 males swarming at a time. In contrast, flight dynamics parameters were first computed for each mosquito in the group, and then averaged over each group to form a replicate. For females exposed to male sound, a total of 10 to 12 replicates per sound level and species were tested (against a total of 9 to 10 replicates per sound level and species for males exposed to female sound in the reciprocal test). Each replicate was performed on a different day.

The sound and video response parameters were analysed using a Bayesian Linear-Mixed Model (*blmer* function, lme4 package, R). Stimulus sound levels and species were considered fixed effects and days, for which replicates were performed, were considered random effects. Sexes were considered separately. Stepwise removal of terms was used for model selection, followed by likelihood ratio tests. Term removals that significantly reduced explanatory power (*p*<0.05) were retained in the minimal adequate model *(33)*. An additional one-sample t-test (with BF-correction for multiple comparisons) was performed independently for each distribution to measure the significance of the mean to 0, which is the “no response” reference. All analyses were performed using R (version 3.5.3).

## Results

### Typical sound level of a 70-male swarm and species-specific cues

In the soundproof chamber with semi-absorbent walls (reverberation time of 0.05 s in the first-harmonic frequency band), the first-harmonic sound pressure level (‘SPL’: root-mean-square SPL ref 20 μPa; see Supplemental Methods section ‘Sound pressure level’) of a station-keeping swarm of ~70 male *An. coluzzii* was 20±3 dB at a distance of 0.9 m from the microphone to the swarm centre, which was 0.6 m high (Figure 1).

The sound of a swarm is composed of the flight sound of individual males. As they probably cannot synchronize the phase of their wingbeats and since the sound of a swarm from a distance is relatively steady over time (for a swarm composed of > tens of individuals), the only species-specific sound cues of a swarm, if any, would come from the frequency content (i.e. not from specific sound phases or time-changing patterns). Sound S1 and Sound S2 are the male sound stimuli used for playback for each of the two species, respectively (before any filtering; Figure S4). Figure S4C shows the strong similarity between the sound spectra of the swarm stimuli of the two species, *An. coluzzii* and *An. gambiae s.s*.: the relative second and third harmonic amplitudes were the same; the fourth-harmonic amplitudes differed, but their respective frequencies were both far above mosquito audibility *(3)*; the mean swarm wingbeat frequencies differed slightly by 21 Hz (857 Hz for *An. coluzzii* and 836 Hz for *An. gambiae s.s.)*, but with a large overlap of 47 Hz of the harmonic peak bandwidth at −3 dB. Note that the 30-male *An. gambiae* swarm sound-level was increased to be the same as that of 70-male *An. coluzzii* swarm, as shown in Table S2, by using the *An. coluzzii* first-harmonic amplitude as a normalization factor (see Supplementary Methods section ‘Sound stimuli’).

### Females do not respond to male swarm-sound, at least up to 48 dB SPL

We played-back the sound of male swarms to a group of 1-5 swarming *An. coluzzii* females at four different sound levels (Table S2) and we tested whether the females responded to the sound stimulus by changing their wingbeat frequency or flight trajectory dynamics (*n*=10 to 12 replicates per sound level, depending on the sound stimulus). The reciprocal was done with 1-6 swarming males exposed to the sound of swarming females, as a control (*n*=9 to 10 replicates, depending on the sound stimulus). Sound S3 and Sound S4 are the female-swarm sounds of the two species, respectively (before any filtering; Figure S4).

Figure 2 B shows the distribution of positions (in three dimensions), linear speed, angular speed and mean wingbeat frequencies produced by groups of 1-5 females or 1-6 males, before, during and after exposure to the loudest opposite-sex sound stimuli (48±3 dB SPL). For each replicate and for each stimulus sound level, we measured the difference between the median wingbeat frequency over the first second of the sound stimulus and during the second before the sound stimulus to monitor response at the sound onset. We did the same for the angular speed.

Our results (Figure 4 A) show that free-flying females do not respond to the sound stimuli by changing their median angular speed with respect to the tested SPLs (LRT, χ_1_^2^=3.81, *p*=0.051) and no angular speed distributions were significantly different from the intercept (i.e. no angular speed change) including the loudest 48 dB SPL distribution (one-sample *t*(22)=1.04, BH-corrected *p*=0.31, mean=0.3 rad/s for a mean angular speed of 4.6 rad/s in absence of sound stimulus). Similarly, females do not respond by changing their median wingbeat frequency with respect to SPLs (LRT, χ_1_^2^=3.19, *p*=0.074) or with respect to the intercept, including the loudest 48 dB SPL distribution (one-sample *t*(22)=0.48, BH-corrected *p*=0.64, mean=1 Hz for a mean wingbeat frequency of 487 Hz in absence of sound stimulus).

**Figure 4.**
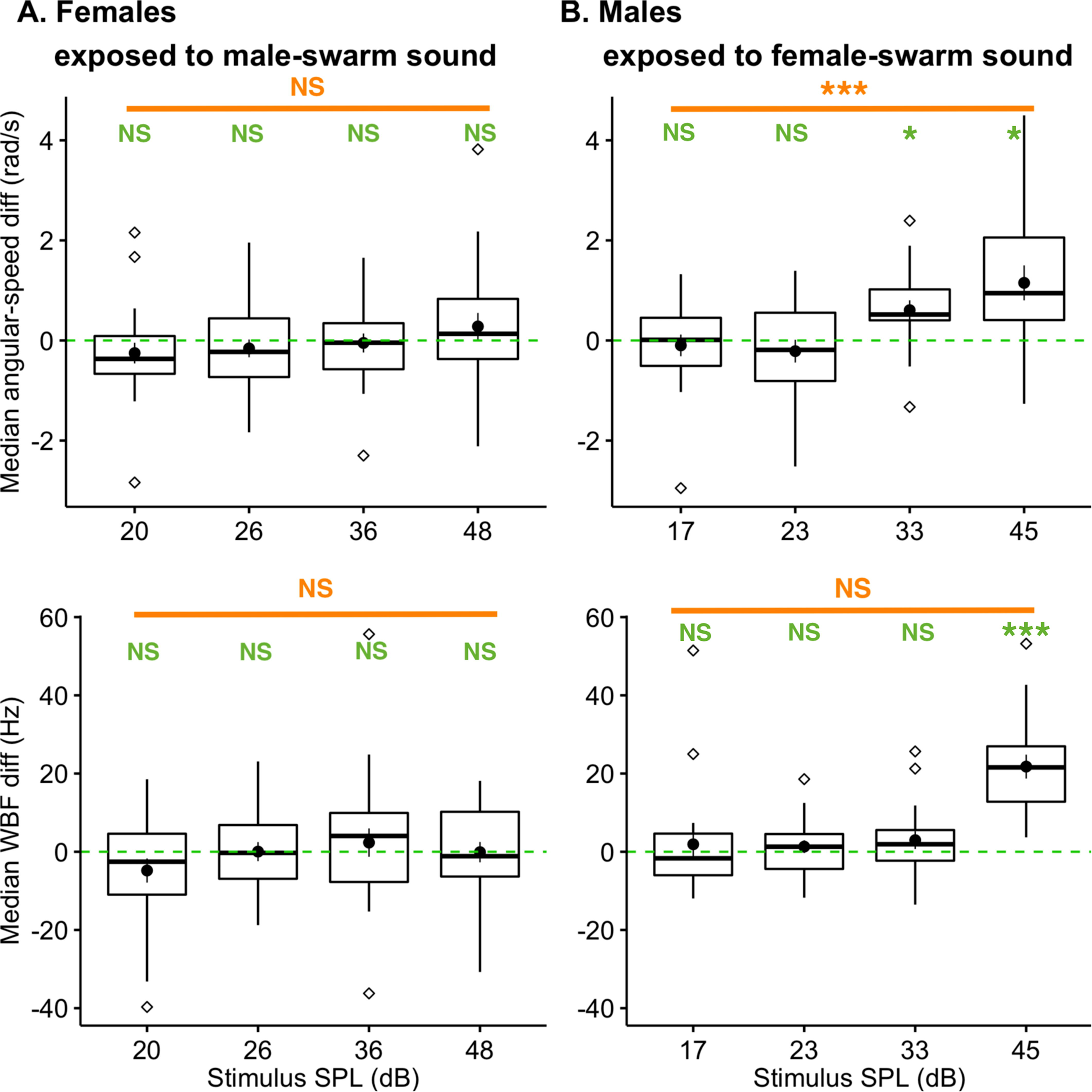
Results of behavioural experiment. Acoustic parameters (e.g. here median wingbeat frequency difference over 1 s, bottom row) and flight parameters (e.g. here median angular speed difference over 1 s, top row) were extracted from flight tracks and wing-flapping sound for statistical analyses of female data (left column) and male data (right column). ‘Zero’ (green dashed line) indicates no difference in the metric before and during the sound stimulus. Boxplots of the parameters show the median, 2^nd^ and 3^rd^ quartiles. Outliers shown as diamond shapes are outside the interval [*Q1* – 1.5 * *IQD*, *Q3* +1.5 * *IQD*] which shows as whiskers (*Q1* = first quartile; *Q3* = third quartile and *IQD* = interquartile distance). Black disk in each distribution shows mean and standard error. Two independent types of statistical tests were performed. Stepwise removal of terms was used for model selection, followed by LRT (likelihood ratio tests, see orange annotation for each of the four plots). An additional one-sample t-test with BF-correction for multiple comparisons (see green annotations above each boxplot) was performed independently for each distribution to measure significance of the mean to zero value (dashed green lines). **(A)** Female *An. coluzzii* responses to *An. gambiae s.l*. male-swarm sounds at four SPLs. For the parameter related to angular speed and the one related to wingbeat frequency, there was neither an effect of SPL nor a significant difference to the baseline (see the Results section for statistical values). **(B)** Male *An. coluzzii* responses to *An. gambiae s.l*. female-swarm sounds at four SPLs. For the wingbeat frequency and the angular speed parameters, there was a strong effect of the SPL, with a significant one-sample t-test for the 33 dB and/or 45 dB SPL distributions (see the Results section for statistical values).

Males are known to be generally more sensitive to mosquito flight sounds than females *(13, 22, 34, 35)*. Accordingly, males were exposed to swarming female sounds, as an experimental control, to demonstrate the relevance of our protocol for assessing female responses to sound. This reciprocal test of male response to female sound stimuli resulted in a highly significant response (Figure 4 B). Indeed, for males, the effect of SPL was to increase their median angular speed (LRT, χ_1_^2^=16.5, *p*<0.001), and the 33 dB and the 45 dB distributions were highly significantly-different from the intercept (respectively: one-sample *t*(17)=3.09, *p*=0.013, mean=0.6 rad/s; sample *t*(17)=3.30, *p*=0.013, mean=1.2 rad/s; for a mean angular speed of 4.6 rad/s before the sound stimulus). Similarly, the effect of SPL was to increase their median wingbeat frequency (LRT, χ_1_^2^=24.6, *p*<0.001), and the 45 dB distribution was highly significantly-different from the intercept (one-sample *t*(17)=7.11, *p*<0.001, mean=22 Hz for a mean wingbeat frequency of 803 Hz before the sound stimulus). However, there was no effect of the SPL on the median linear velocity, median height or median distance to the speaker (Table S3).

Given the absence of female response to male sound, we decided to increase the number of tested parameters to be certain we did not miss any meaningful variables. Table S3 gives an extra four flight-dynamics parameters tested (linear speed, height and distance to the speaker), calculated at the stimulus onset (1 s) or considering the full stimulus duration (7 s). Benjamini & Hochberg correction of *p*-values for multiple comparisons led to no statistically significant predictors of female response in terms of SPL, but also no effect of species, or SPL and species interaction, as expected by the absence of significant differences in the swarm sound of the two species.

These results support the proposition that a female cannot hear male-swarm sound stimuli below 48 dB SPL, but higher sound levels were not tested. Therefore, the hearing threshold for females could be above 48 dB SPL.

### Females cannot hear a 70-male swarm as a whole before she hears peripheral males

Given that our ~70 male *An. coluzzii* swarm was 20±3 dB at a distance of 0.9 m, we calculated the equivalent distance corresponding to the sound of a 70-male swarm modelled as an acoustical point source, at 48±3 dB SPL, which is the loudest sound level we tested. This distance is equal to 0.04±0.01 m if considering the far-field assumption (under-estimated at this distance), and anyway < 0.15 cm if not considering the far-field assumption (Table S1; see Supplementary Methods section ‘Relationship between particle-velocity and pressure levels’ for discussion related to reproducing a sound-source outside the far-field range). This distance is less than the swarm radius of the 70-male swarm which we recorded in the laboratory (0.2 m). As a consequence, if a female cannot hear this sound level, then a female flying close to a real 70-male swarm will hear the peripheral male nearest the female before she would be able to hear the swarm as a whole. Indeed, at this distance, a peripheral male near the female will produce a sound that is louder than that of the rest of the swarm as a whole because of the rapid increase in particle velocity in the vicinity of a mosquito. Therefore, we conclude that a female cannot hear a 70-male swarm as a whole until she is within its boundary.

### How far away is a >48 dB SPL swarm of a given number of males?

Using another acoustic prediction formula relating the sound level to the number of mosquitoes (see Supplementary Methods section ‘Formula relating hearing distance and number of individuals in the swarm’), Figure 5 shows the female hearing ranges as a function of distance to the swarm and number of males in the swarm. The previous findings (no response up to 48 dB SPL) allow us to split the 2-D plot into the ‘no-response’ area (red) and the ‘unknown’ area (white). The hearing distance threshold may stand somewhere in the white area.

**Figure 5.**
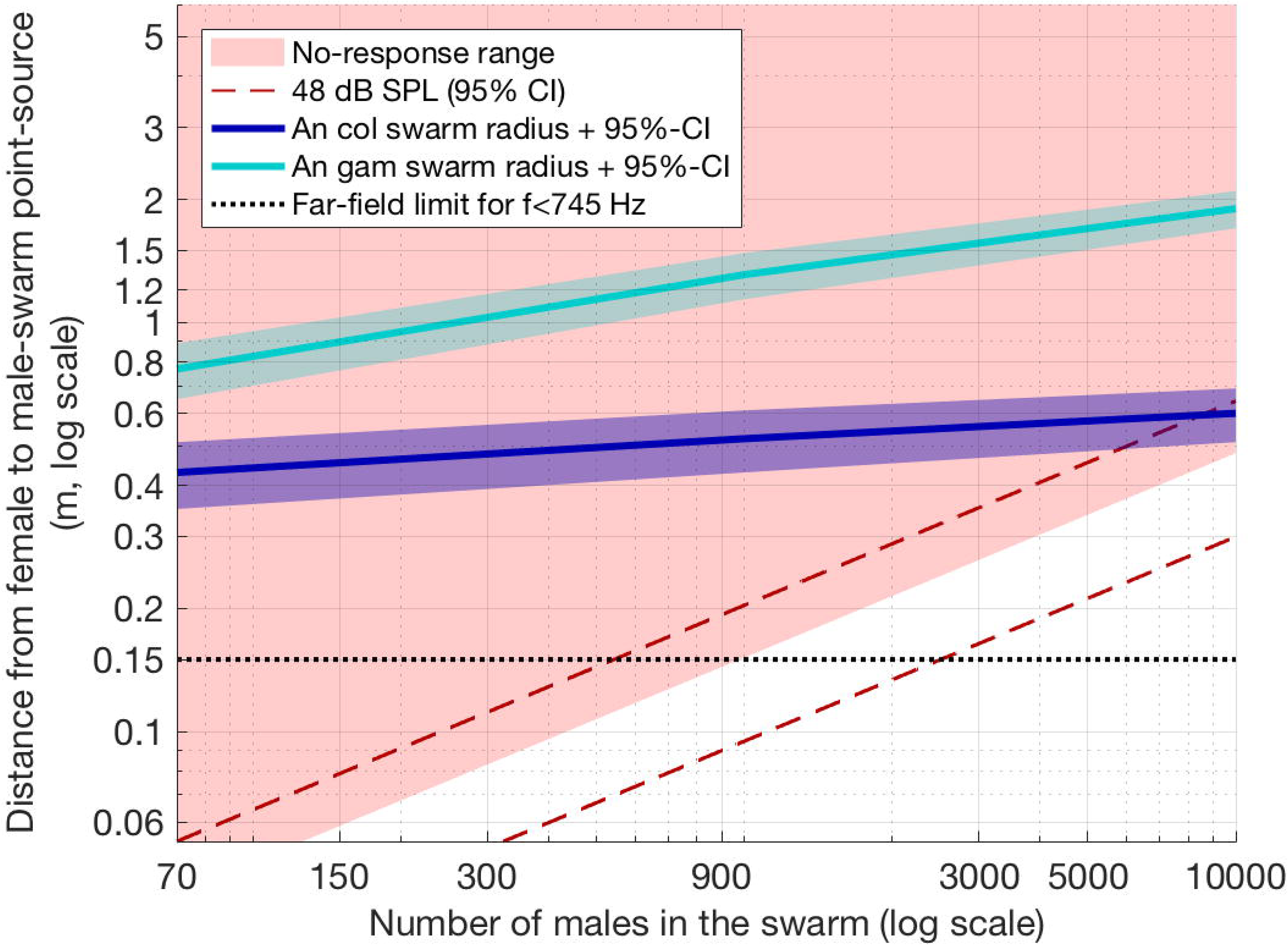
Estimated no-response range and swarm radius as a function of the number of males in the swarm. Red area indicates the minimal non-response range of a female to male swarm sound for both species, as a function of the number of males in a given swarm (X-axis) and the distance to the swarm centre (Y-axis). These areas are based on our behavioural results showing a non-response for stimuli equal or quieter than 48 dB SPL stimulus (red-to-white boundary), with 95% confidence interval (dashed lines). The swarm is assumed to be a point source in the model and only the far-field component of the particle velocity is considered (see Methods section ‘Acoustic assumptions’): above 0.15 cm (black dotted line), the near-field component of the particle velocity is negligible (< 1 dB); below 15 cm the smaller the distance, the less linear the relationship between distance and number of males is (i.e. the line forming the boundary between the two coloured area should be higher than shown on this graph). The light and dark blue lines, along with their 95% CI, represent the estimated mean swarm radius of 95% of *An coluzzii* and *An. gambiae s.s*. swarming males (see Figure S2).

For illustration, a point-source swarm of 1,000 males of ≥ 48 dB SPL can be expected to be at a distance of ≤ 0.15±0.05 m and a 10,000-male swarm of same SPL at a distance of ≤ 0.5±0.2 m. Table S1 incorporates all the acoustic sound levels related to the listening distance and to five orders of magnitude in the number of males. This results in two statements: 1) there is no chance that a female can hear a swarm at a distance less than 0.8 m from its centre for a number of males up to 10,000, which we consider the upper limit (see Supplementary material section ‘Number of males in swarms’) and 2) there is no chance for a female to hear a swarm before reaching the peripheral males if the swarm is less than a couple of thousands of males. In order to conclude with more confidence for the largest swarms, we need to estimate their dimension, which we did by using data from *An. gambiae s.l*. swarms in the field *(18)*.

### Females cannot hear swarms before entering them considering their dimension

An extrapolation of the Manoukis *et al* data *(18)* (see Supplementary information ‘Swarm radius as a function of number of males’) shows that our estimate of *An. coluzzii* swarm radius is consistent with observations of swarms with thousands of males which are usually < 1 m in radius *(36)*. For *An. coluzzii*, the predicted mean swarm radius is 0.5±0.1 m for 95% of 1,000 swarming males (0.20±0.05 m for 50% of them) and 0.6±0.1 m for 95% of 10,000 males (0.21±0.05 m for 50% of them), representing a steep increase in density of swarming males, especially in the swarm centre (Figure S2). The swarm radius of an *An. gambiae s.s*. swarms is slightly larger for small swarms, but the predicted radius for large swarms is much larger (Figure S2). Figure 5 shows the 95%-male swarm radius of both species, which are in the ‘no-hearing’ area, showing that *An. coluzzii* females cannot hear male swarms before entering them, even considering the loudest swarms of 10,000 mosquitoes.

## Discussion

### Hearing sensitivity of *An. coluzzii* females and males

Previous studies estimated the hearing threshold of tethered *An. gambiae s.l*. females was in the range 44-52 dB (particle velocity of 14±6 μm.s^−1^, *n*=5) and tethered *Aedes aegypti* females around 55 dB SPL (*n*=10) by monitoring the activity of the Johnston’s organ nerve *(4, 16)*. In the present study, we did not elicit a behavioural response in free-flying *An. coluzzii* females with 48±3 dB SPL. For free-flying *An. coluzzii* males, we found a significant response at 33±3 dB SPL, and no response at 23±3 dB, indicating that their hearing threshold are likely to be < 33±3 dB. This is a lower threshold than reported values for tethered male *An. gambiae s.l*. (18±6 μm.s^−1^, i.e. 38-39 dB SPL for the SD range in the far-field, *n*=5) from recordings of the Johnston’s organ nerve with the antenna fibrillae extended *(4)*, but similar results to tethered male *Culex pipiens pipiens* (32.0±4.4 dB sound particle-velocity level, n=74, equivalent to 32.0±4.4 dB SPL in the far-field) *(37)*.

To our knowledge, this study is the first report of sound sensitivity based on behavioural responses in free-flying mosquitoes. The higher sensitivity in *An. gambiae s.l*. males than those reported in electrophysiological studies can be explained by active hearing *(7, 38)*, which could be triggered only by using natural behaviours (i.e. free-flight and mating behaviour). Furthermore, we played-back the sound of a large group of swarming females (i.e. wide band tone) to test male sensitivity, which does not occur in the field. Accordingly, we still expect a greater sensitivity for free-flying males exposed to single-female sound (i.e. sharp-band tone corresponding to the sound of a single female), as noted previously *(12)*. In the case of females, we expected to trigger a response at lower values than previously reported, i.e. 45 dB SPL or lower in our case, because we used a natural behaviour. More investigations have to be carried out to relate the female’s electrophysiological and behavioural responses.

### Long-range hearing does not contribute to conspecific mating

First, species-specific cues of swarm sound were found to be weak (Figure S4). Second, our behavioural assay did not show any species-specific responses in *An. coluzzii* females to the swarming sound of *An. coluzzii* or *An. gambiae s.s*. males. Third, following our results, we can reject the idea that females use the sound emanating from a swarm to determine whether to avoid entering the swarm of the wrong species, or to join the swarm of the same species, because the female will not hear the swarm before she comes into close proximity of numerous males at the periphery of the swarm.

Swarm localization by females is much more likely to occur via responses to the same environmental cues as their male counterparts, enhancing the likelihood of encountering conspecific males. It is possible that long-range cues are not necessary for the female to arrive at a swarm site. In that case, females may use the close-range sound of a chasing male to avoid being inseminated by the wrong species *(4)*, however, investigations on long-range cues such as vision *(29)* or olfaction *(39, 40*) should be pursued in future research.

### Long-range hearing is unlikely in inter-mosquito communication

To our knowledge, male swarms are the only likely candidate source of sound which is loud enough and fits the tuning of the mosquito organs to enable inter-mosquito acoustic communication at long-range. This study presents data that rejects the hypothesis that *An. coluzzii* females can hear a male swarm before entering it. It is also unlikely that a male hears a male swarm at long-range because, although males are more sensitive to sound than females *(13)*, their hearing organ is not tuned to male wingbeat frequencies. Finally, as we chose a mosquito species which produces large and loud swarms, we can claim that long-range interspecific acoustic-communication in mosquitoes is unlikely to occur before the female mosquito enters a swarm.

However, this study does not eliminate the hypothesis that long-range hearing can be used for host location *(23, 41)* or for predator avoidance *(22)*, providing the host/predator sound is loud enough and tuned to mosquito hearing.

### Limitations of the experimental design

The main limitation of our experimental paradigm is that we used swarming females to test their response to male-swarm sound (see Methods section ‘Experiment paradigm’). This means that when females were exposed to the swarm sound in the laboratory, they were already flying above a swarm marker, while in the field they would have been responding to environmental stimuli leading them to a swarm marker. This may have induced females to continue swarming over the marker without altering their behaviour when male sound was played-back, effectively waiting for males to approach the marker. However, we monitored all the likely flight variables (flight velocities, positions and wingbeat frequency changes), so it is unlikely that we overlooked any female reactions to sound and unlikely that females would not respond if they could hear a male sound.

Instead of a highly complex model of the swarm acoustic consisting in individual dipole sources, we chose a simple model of the swarm by modelling it as a single monopole sound object. While well justified at long-range (see Methods section), this has limitations in terms of sound spatiality and particle velocity field when approaching the swarm at closer ranges. However, our model is conservative as respect to our results because 1) dipole SVL at short range decreases with distance quicker than if considering monopole source *(15)* and 2) if the swarm is not loud enough when considering all male sounds virtually packed in a point source, it also won’t be loud enough to trigger a response in the case of a normal spatial distribution of males around the swarm centre. Then, the first sound eliciting a response will be from a single peripheral male which will trigger the mating chase, but one-to-one interactions were not within the scope of our study.

## Supporting information

Sound S1

Sound S2

Sound S3

Sound S4

## Acknowledgments

David Sanou (swarm location in Bama), Natalie Morley (insect rearing), Dr Paul Luizard (proof-reading of some acoustic equations), Stephen Young (discussion about statistics), Greg Smith from IAC Acoustics (tips on room acoustic). Opinions, findings, conclusions, and recommendations expressed in this publication are those of the author and do not necessarily reflect the views of the USDA. The USDA is an equal opportunity provider and employer.

## Author contributions

Conceptualization LF, GG, OR; Methodology LF, GG; Software LF; Formal Analysis LF; Investigation LF; Resources GG, OR, LF, NM; Data Curation LF; Writing – Original Draft LF, GG; Writing – Review & Editing OR, NM; Visualization LF; Supervision GG, OR; Funding Acquisition OR, GG, LF.

## Data and materials availability

Custom audio-video code for parameter-extraction, audio-video synchronization (Matlab files), custom statistics code for data analysis and figure plot (R files), and dataset (Text file) are available at http://dx.doi.org/10.17632/hn3nv7wxpk.2. Raw sound files and tracked flight dataset are available on request.

