## Supplementary Materials for "Mosquito sound communication: are male swarms loud enough to attract females?"

Mosquito sound communication: assessment of ecologically relevant ranges

### Supplementary Methods

#### Environmental conditions in soundproof chamber

The swarming arena was designed to include the key environmental conditions and sensory cues known to control mating and swarming flight in the field. A large mosquito bed-net enclosure (NATURO, *L* x *W* x *H* = 1.8 m x 1.7 m x 2 m) filling most of the soundproof chamber (Figure 1) enabled mosquitoes to fly freely in a volume 100 times greater than the volume covered by the swarming space.

***Light and visual cues.*** Lighting was provided by an artificial-sunlight system to imitate natural daylight, sunrise and sunset (LEDs 5630, HMCO FLEXIBLE dimmer, and PLeD software, custom-built). Lamps were arranged to mimic sunset lighting; a sharp horizon ~40 cm above the floor on one side of the room provided a 'sunset' feature and a gradually decreasing light intensity with increasing height above the floor (Figure 1). Ceiling lights were dimmed over 30 min, while the horizon lights started to dim 10 min before the ceiling light turned off, whereupon the light intensity decreased gradually over 10 min and finally remained constant for 1 h to provide a constant very dim light intensity that favored prolonged swarming flight during the experiments. The visually conspicuous matt-black swarm marker triggered swarming behaviour. The marker consisted of a circle of matt-black paper (Ø=30 cm), placed > 30 cm away from the closest netting. The location and height of the swarm marker was arranged according to the swarming behaviour of each species in order to induce swarming flight at the same location in the room for the two species: therefore, the swarm marker for *An. gambiae s.s.* was raised by 6 cm and moved 0.8 m horizontally in the opposite direction of the dusk light, compared to the position of the *An. coluzzii* swarm marker (Figure 1), as previously reported *(S1)*.

***Temperature monitoring.*** The temperature was monitored by type-T thermocouples (IEC 584 Class 1, Omega) associated with a temperature logger (HH506RA, Omega) with a measurement accuracy error of ±0.9 °C. The chosen thermocouple was located on a room wall at a height of 85 cm from the floor. The four recordings of the reference sound stimuli (two species, two sexes) were recorded at 28.0 °C. The mean temperature and standard deviation of the behavioural assays were 28.0±0.3 °C.

#### Recording environment

***Soundproof chamber.*** All experiments were conducted in a soundproof chamber to limit interference from external sounds. The chamber consisted of double-skin soundproof walls, ceiling and floor (*L* x *W* x *H* = 2.7 m x 1.9 m x 2.3 m), with carpet on the floor, semi-absorbent internal walls/ceiling and a layer of white cotton cloth covering all surfaces, producing a reverberation time ≤ 0.07 s for frequencies above 200 Hz (measurements conducted in empty room by IAC Acoustics, manufacturers). Figure S5C displays the sound level per octave band when the soundproof chamber was silent (dashed lines). At low frequencies (<176 Hz), the sound pressure level (SPL) was ≥ 25 dB (ref 20 µPa). Between 176 Hz (lower limit of the 250-Hz octave band) and 1.4 kHz (upper limit of the 1-kHz octave band), i.e. the frequency range within the *An. coluzzii* mosquito's response is the highest *(S2)*, the SPL was < 14 dB, which is 8 dB less than the quietest sound stimulus used in the study.

***Sound monitoring.*** The wingbeats (aka, 'flight tones') of mosquitoes in the laboratory were recorded with a weatherproof microphone (Sennheiser MKH60; RF-condenser; super-cardioid polar pattern at 0.5-1 kHz, with amplitude decrease of > 15 dB beyond 90° from the microphone head) directed toward the swarm location. The microphone was located at a distance of 0.9 m from the centre of the swarm area (Figure 1) and plugged into a Scarlett 18i8 audio interface on a Windows7 computer running Pro Tools First 12.8 (Avid Technology, Inc).

***Flight track recording.*** The 3D flight trajectories of mosquitoes were captured at a sampling rate of 50 Hz with Trackit software (SciTrackS GmbH, Switzerland, *(S3)*). Two video cameras (Basler, ace A640-120gm) were fitted with wide-angle lenses (Computar, T3Z3510CS, 1/3" 3.5-10.5mm f1.0 Varifocal, Manual Iris) to maximize 3D volume of video-tracking and infra-red bandpass filters (>840nm, Instrument Plastics Ltd UK). Ten IR lights (Raytec RM25-F-120 RAYMAX 25 FUSION) enabled the tracking system to detect flying mosquitoes as silhouettes against an IR-illuminated white back-wall made of thin cotton cloth (Figure 1). The dual IR/white lighting system enabled constant bright IR light (invisible to mosquitoes) for video-tracking flying mosquitoes. The 3D-flight trajectories were smoothed using a cubic spline interpolation at a sampling frequency of 200 Hz on Matlab (version R2017a).

***Field recording.*** Preliminary recordings of the flight sound of wild male *An. coluzzii* swarms in the area where our colony originated from (village VK5, Bama, Burkina Faso, 11°23'17.5"N 4°24'27.0"W, October 2017) were used to study the signal-to-noise ratio of swarm sound in the field against local background noise. The swarm was spherical (~1 m diameter), centred ~3 m above the ground and was not apparently disturbed by our presence and produced sound at acceptable levels for recording. The swarm consisted of several thousands of *An. coluzzii* (estimated by eye, by LF, OR and experienced technical staff from the IRSS). The swarm's sound was recorded from various positions and distances; from ~0.2 to 3 m away. The recordings were produced with an RF-condenser microphone (MKH 60 P48) plugged into a Scarlett Solo sound card, captured via Audacity (v2.2.1) software on a MacOS operating system (v10.12.6).

**Choice of species as test subjects and for production of sound stimuli.**

We had no difficulty in triggering robust swarming behaviour in *An. coluzzii* males and females and in *An. gambiae s.s.* males, but it was difficult to obtain consistent results with *An. gambiae s.s.* females. For this reason, we focused on the response of *An. coluzzii* to sound stimuli of each species. Female responses to male flight sound were absent, therefore, we conducted the reciprocal experiment with *An. coluzzii* males exposed to female-swarm sound, which confirmed that the experimental protocol was valid (male responsiveness to female-swarm sound was robust). Although it was more difficult to induce *An. gambiae s.s.* females to swarm, we recorded the sound of a swarm composed of 4 females at a time, *versus* 30 females at a time for *An. coluzzii*.

#### Generation of sound stimuli

***Recording of the reference sound-stimuli.*** Swarms of *An. coluzzii* females or males, and *An. gambiae s.s.* females or males were recorded in the soundproof chamber (Figure 1 and Figure S4). About 300 x 4-7 days-old males or 1-4 x 2-6 days-old females were released in the swarming arena two days before the experiment to acclimatize before their flight sounds were recorded. The standard environmental conditions in the room were: 12h:12h light:dark cycle with a 1h artificial dawn/dusk transition in light intensity, 21-28°C (warmed up to 28°C before the mosquitoes flew) and ~60-75% RH. One recorded 7s sequence was selected for each sex/species, which began ~10 min after the first male/female started to swarm (Sound S1: *An. coluzzii* male swarm; Sound S2: *An. gambiae s.s.* male swarm; Sound S3: *An. coluzzii* female swarm; Sound S3: *An. gambiae s.s.* female swarm). The swarms were composed of 30-70 individuals (except for the *An. gambiae s.s.* female swarm: 4 individuals) flying in loops 0.3 m above the floor level with a horizontal diameter of 0.2 m. The start and the end of the recorded-sound amplitude were multiplied by a fading window of 1 s to avoid creating sound artefacts due to the signal truncation, and to make the stimulus more natural, i.e. mimicking the male swarm sound amplitude which continuously increases when the female gets closer to the swarm.

***Reference sound-stimulus gain***. For each sex, the *An. coluzzii* swarm was the reference, and to balance the different number of individuals in the swarms of the two species, the *An. gambiae s.s.* swarm sound level was adjusted to that of *An. coluzzii* (based on the 50Hz-smoothed spectrum peak of the first harmonic, which is known to be important in mosquito hearing *(S2)*). We took advantage of the high numbers of *An. coluzzii* mosquitoes that swarmed (70 males and 30 females), which we did not achieve with *An. gambiae s.s.* (30 males and 4 females), even though it meant adjusting the sound level of *An. gambiae s.s.* sounds stimuli (see Figure S4).
To playback the stimuli at natural sound levels, we first played them back in the same room and at a distance to a speaker (Genelec 8010A) identical to the distance between the swarm and the microphone. Second, the gain was set to ensure the same relative sound pressure level was used as during the reference swarm recording (based on the first harmonic amplitude peak from a 50Hz-smoothed spectrum, Figure S4, and using the same software and hardware settings).
Figure S4 gives the sound spectrum of the swarm sounds used in the assays. The female harmonics from three times the fundamental frequency were filtered out in order to free some spectral space for male wing beat tracking, which does not change the response to the sound stimuli since these higher harmonics are unlikely to be heard by these mosquitoes *(S2)*.

***Generating the different sound levels.*** In addition to the natural sound level of the reference sound stimulus (i.e., 70-male swarm or 30-female swarm 0.9 m away), we generated three more stimuli for each species and each sex, to test the efficacy of sound levels over the range of the response possibilities, using Matlab (R2017a, The Mathworks Inc, Natick, USA) at a sample rate of 8 kHz / 24 bits. The additional gains applied to the natural-sound-level reference stimuli were computed using a criterion based on the maximum value of the first harmonic on a 50-Hz-smoothed sound spectrum: +6.0 dB, +16 dB and +28 dB compared to the reference sound stimuli (see Table S2 for measured SPL of each stimulus).

A high-pass filter was added to remove the electrical noise below the first harmonic (without removing any frequency component of the swarm sound). The eight stimulus sounds (two species x four sound levels) were combined sequentially with a 10 s silence interval. Ten sequences were generated, each containing the four sounds ordered randomly.

***Sound stimulus playback.*** Recorded mosquito sounds were played-back from a speaker (Genelec 8010A) with its membrane located 57 cm above the floor, 15 cm from the back wall, and 0.9 m from the swarm marker (Figure 1). Both microphone and speaker were plugged into a Scarlett 18i8 sound card running pro Tools First and Audacity on Windows 7.

**Response Parameters**

***Wingbeat parameter extraction from flight sound.*** Wingbeat frequency was tracked every 40 ms using a Fast Fourier Transform algorithm (256-ms FFT, Hanning-windowed). Since females and males do not have the same wingbeat frequencies and we always played-back opposite-sex sound stimuli to individuals, we had to operate differently for each sex. For females, their fundamental wingbeat frequencies were tracked between 370 Hz and 660 Hz (given that the mean female wingbeat frequency was 487 Hz) to avoid overlap with played-back wingbeat harmonics of swarming males (female wingbeat frequencies were always lower). For males, only the two first harmonics of female sound stimuli were played-back as explained above and then the male's third harmonic (3 x fundamental frequency) was tracked between 2190 and 2920 Hz (given that the mean male's third harmonic without sound stimulus was 2409 Hz), since it is the lowest harmonic that does not overlap with the sound stimulus (example of spectrogram in Figure 2 C). When several wingbeat frequencies were tracked due to the presence of several mosquitoes over the swarming marker, their wingbeat frequencies were averaged. Male harmonics were divided by three to get the fundamental frequency. Finally, a 3-point median filter was applied over time to reduce wingbeat tracking error. Figure 2 C gives an example of detected wingbeat frequencies of females and males (Figure S3 gives the superimposition of all tracks for a same stimulus) while Figure 2 B shows the distribution of the detected wingbeat frequency over time for all recordings associated to the loudest stimuli.

***Speed parameter extraction from tracked flight trajectory.*** The criteria used to include a tracked flight in the data analysis was that the mosquito was swarming over the marker for at least 1 s before and after the sound stimulus onset. Linear speed at time index *n* was calculated as the square root of the sum of the three square velocity components provided by the Trackit software, and the angular speed was computed as $avel=\Delta\theta/\Delta t$, where $\Delta t = t_n - t_{n+1}$ is the duration between two consecutive time indexes *n* and *n*+1, and $\Delta\theta$ is the turn angle defined as:

$$\Delta\theta = cos^{-1}\frac{v_n . v_{n+1}}{|v_n| . |v_{n+1}|} \qquad (1)$$

where $v_n$ is the three-dimensional linear velocity vector of the mosquito at time index n and $|v_n|$ is its magnitude.

***Sound and video synchronization.*** To synchronize sound and video data, a custom-made 'clapper-board' simultaneously switched off an IR led and a 3900-Hz bip sound (which cannot be heard by this species complex *(S2)*). The IR light was located on the edge of the field of view where no mosquito was expected to swarm. The IR light was tracked with the software Trackit every 2 ms when the light was switched off (i.e. creating a dark silhouette) simultaneously with the sound and was automatically detected on Matlab. The 10-s bip sound was played-back before and after each stimulus sequence and manually switched off along with the IR light. The bip 'offsets' were detected manually on an 8 ms-window spectrogram. Cumulative errors over time were controlled by using the 'offset' time before and after the stimulus sequence. Overall, the synchronization uncertainty was ±8 ms.

#### Sound pressure level (SPL)

***Measurement.*** Stimulus SPLs were measured at the females' swarming position with a sound metre (Casella, CEL633C1, Class 1) set as follows: reference pressure of 20 μPa; no octave weighting (i.e. dB Z); slow octave time-constant (IEC 61672-1: 2002); octave and third-octave bands; calibrated twice a day (CEL-120/1, Class 1, at 94 dB / 1 kHz) before and after each measurement. The minimum and maximum sound level values within each stimulus duration were used to compute the mean and error of each measurement (Table S2). The speaker and the software/soundcard gains were set to be the same as during the behavioural experiment.

All SPLs reported in the paper take into account only the frequency bands that are audible by mosquitoes, i.e. mostly the first-harmonic of the opposite sex *(S2)*. They were calculated as follows: $10log_{10}(10^{0.1L_{B1}} + 10^{0.1L_{B2}})$ where $L_{B1}$ and $L_{B2}$ are SPL measurements in frequency bands $B1$ and $B2$; $B1$ and $B2$ are the third-octave bands nearest the wingbeat frequency of the first-harmonics, i.e. 800 Hz and 1000 Hz for males and 500 Hz and 630 Hz for females (Table S2; and Figure S5 for all third-octave values). This method enabled us to compare our sound stimulus levels to pure sounds used in previous studies on hearing sensitivity and is closer to what mosquitoes actually hear.

***Estimate of SPL errors at mosquito's location.*** Three types of SPL errors were taken into account. The first is related to the time variation of the sound stimulus levels which were between ±0.3 dB and ±1 dB, depending on the stimulus considering maximum error (see Figure S4 for an example of stimulus RMS-pressure-level along time).

The second source of error is related to acoustical interferences caused by room boundaries. Up to this point, we have considered a free-field acoustic-propagation hypothesis to simplify the problem. In a room, however, sound level can decrease (destructive interference) or increase (constructive interference) independently of the distance to the speaker. This effect was reduced by the semi-absorbent walls of the room but was still present because the room was not an anechoic chamber. Boundary-induced 'comb filtering*'* was reduced by locating the speaker close to the wall, but acoustic room modes were still present. We played-back the *An. coluzzii* male and female swarm stimulus and measured the sound level in a 0.2 m diameter sphere around the expected swarm centre. The maximum error was about ±1 dB for the female sound stimulus and ±2 dB for male swarm stimulus. We ignored any reverberation effect, as we estimated its effect to < 0.4 dB at 800 Hz, using an acoustic room model of the ratio of direct and reverberant sound, given the reverberation times of the room provided by the soundproof chamber designer (IAC Acoustics Ltd).

The last type of measurement uncertainty arises when the estimated sound level should be estimated from the mosquito position. SPLs were measured at the expected centre of the station-keeping swarm-flight of the mosquito. However, the distance between the female and the speaker varies between 72 and 113 cm (95%-CI, Figure 2 A) due to the females' swarming-flight pattern and sound level changes, accordingly. We computed this error by considering the fluctuating distance between the female mosquito and the speaker using equation 4.

Finally, using standard uncertainty-propagation theory, we calculated the total error of sound pressure level $L_i$ at the location of the female exposed to male sound, resulting in a total error of ±3 dB SPL for the SPL. This error is considered to be conservative (at least 95%-CI) and were used to interpret the results of the experiments. For errors related to the difference between what we measured (sound pressure) and what mosquitoes detect (particle velocity), see main-text Method section 'Physical sound quantities produced by a speaker and sensed by mosquitoes'.

**Acoustic formulae**

***Relationship between particle-velocity and pressure levels.*** We monitored the sound level of swarms by recording the sound pressure level (SPL), while mosquito hearing organs are sensitive to particle velocity levels (SVL) *(S4)*. These two quantities are equal only far from the sound source, so it is important to understand how they are related to estimate the error when we are dealing with sources close to the receiver.
The root-mean square value (RMS) particle velocity $v_{RMS}$ and the RMS sound pressure $p_{RMS}$ can be related as follows for a point source radiating spherically (air impedance $Z_{air}$(28°C) = 408 N.s.m$^{-3}$; sound speed $c$(28°C)=S118 m.s$^{-1}$) *(S5)*:

$$v_{RMS}(r) = \frac{p_{RMS}(r)}{Z_{air}} \sqrt{1 + \left(\frac{c}{2\pi f r}\right)^2} \qquad (2)$$

The SPL $L_p \stackrel{\text{def}}{=} 20log_{10}(p_{RMS}/p_0)$ and the associated particle-velocity level $L_v = 20log_{10}(v_{RMS} Z_{air}/p_0)$ (sea-level RMS atmospheric pressure $p_0$ = 2.0 10$^{-5}$ Pa) can be calculated as follows:

$$L_v(r) = L_p(r) + 10\ log_{10}\left(1 + \left(\frac{c}{2\pi f r}\right)^2\right) \qquad (3)$$

Therefore, SVL and SPL are equal when $r$ is large. In our case, considering the male swarm sound stimulus does not have any frequency components below $f$ = 745 Hz (the smallest frequency value of the group of first harmonics of the swarming males at -12 dB below the peak at 857 Hz, Figure S4), then we can calculate that for $r$ > 15 cm, $L_v(r) = L_p(r)$ with an error less than 1 dB.

Table S1 gives the SPL of each stimulus, which is equal to the particle-velocity level for distances from the sound-source < 15 cm. Below 15 cm, the smaller the distance to the sound-source, the greater SVL is, compared to the SPL. At 4 cm from the sound source, the SVL is 8 dB higher. When the difference between the SPL and the particle-velocity is greater than 1 dB, the particle-velocity level is added along the distance to the sound-source in Table S1.

***Formulae relating sound level and distance.*** In order to estimate the distance over which a female could hear a given-size swarm with a given number of swarming males, we are interested in determining the equivalent distance $r_i$ ($i$ being the SPL label) to the virtual sound source (i.e. the played-back male swarm, or sound-source image) knowing the SPL $L_i$ at the female's position at a distance $r_i$ from the virtual swarm, and the SPL $L_{ref}$ at position $r_{ref}$ = 0.9 m known to be the distance to the reference sound stimulus source. The physical sound source is the speaker, at fixed distance $r_{ref}$ from the swarm centre (then from the female ± its movement above the swarming marker). The SPL is set to reproduce a natural swarm sound where its presence is virtually located at a distance $r_i$ from the female (see Figure 3 for a visual illustration).

As a single monopole point spherically radiates in all directions (no sound reflection), the root-mean-square sound pressure $p_{RMS,i}$ is inversely proportional to the distance $r_i$ (i.e. $p_{RMS,i} \propto \frac{1}{r_i}$ ) *(S5)*. Then the SPL difference $\Delta L_i$ can also be expressed as follows:

$$\Delta L_i \stackrel{\text{def}}{=} L_i - L_{ref} = 20log_{10}\left(\frac{r_{ref}}{r_i}\right) \qquad (4)$$

Then from equation 4 we get the distance $r_i$ to the sound-source image as a function of the difference level $\Delta L_i$ and of the known distance $r_{ref}$ from the female's position in relation to the sound-source image of the swarm of the reference stimulus recording:

$$r_i = r_{ref}10^{\frac{-\Delta L_i}{20}} \qquad (5)$$

SPL label *ref* corresponds to the natural sound level of an *An. coluzzii* 70-male swarm at a distance of 0.9 m. The equivalent distances $r_i$ associated with the other sound levels $L_i$, ($I \in \{1,2,3\}$) can be calculated from equation 5: they correspond to the SPLs 20 dB, 26 dB, 36 dB and 48 dB of a point-source 70-male swarm at a distance of 0.9 m, 0.5 m, 15 cm and 4 cm, respectively (Table S1). This calculus assumes that the female is far enough from the swarm so that the swarm dimensions are small enough compared to its distance to the swarm (i.e. 'point-source'). Even if the latest is unrealistic for the smallest distance (4 cm), it helps as a step for modelling larger distances where this issue does not occur anymore (see below).

***Formula relating hearing distance and number of individuals in the swarm.*** Acoustic prediction was needed to cope with large swarms because of a limitation in the number of swarming males to be recorded under controlled conditions. In our experimental space, about 20% of the released *An. coluzzii* males and 10% of the released *An. gambiae s.s.* males swarmed over the swarming spot. A small number of the non-swarming males were flying without station-keeping behaviour in our experimental room space (most of the remaining males were resting). However, the chance of a flying non-swarming mosquito passing in the field of sound of the directional microphone increased with the number of released mosquitoes. Thus, above ~70 swarming males, the number of flying non-swarming males was too high and our sound recording could have been altered by flying males for which the distance to the microphone and their behaviour (i.e. non-swarming flight) could not have been controlled. As a consequence, we decided to use the 70-male swarm in the behavioural experiments, which is the biggest station-keeping swarm we could reliably produce and record in the laboratory.

In order to estimate the results which could have been found with a bigger swarm, we predicted the behavioural assay results performed with a 70-male swarm sound stimulus using an acoustic model of the swarm sound level as a function of its number of individuals and its distance to the female.

Multiplying by $N$ a number of acoustically incoherent sources, such as swarming mosquitoes, increases the SPL by $10log_{10}(N)$ *(S6)*. Let's assume a $N$x70 -male swarm can be modelled as a single point (see section 'Acoustic assumptions for a swarm' above), then the SPL at a fixed distance will be increased by $10log_{10}(N)$ (e.g. 7 dB if $N$=5 or 20 dB if $N$=100) compared to the 70-male swarm.

Then we can compute the virtual distances $r_{i,N\text{x}70}$ of a $N$x70-male swarm with same SPL $L_i$ as a 70-male swarm at distance $r_i$, knowing that the $N$x70-male swarm has a SPL $L_i$ + $10log_{10}(N)$ dB at distance $r_i$, by the following formulae derived from equation 4 (values are presented in Table S1 for a 300, 1,500, 6,000 and 10,000-male swarm):

$$r_{i,N\times 70} = r_i 10^{\frac{-\left(L_i - \left(L_i + 10log_{10}(N)\right)\right)}{20}} = \sqrt{N} r_i \qquad (6)$$

#### Supplementary information

##### Mosquito sounds stand out against ambient noise at least 3 m from the swarm

Relative flight-sound intensities of wild male *An. coluzzii* swarms were measured to characterize the sound profile of typical male swarms in relation to the background sounds of other twilight-active organisms, including humans, near rice fields in the Bama village (VK5), Burkina Faso. We recorded ambient sound at ~ 1 m from a swarm consisting of several thousand male *An. coluzzii* around sunset. The recording included background noises from insects, birds, mammals, human speech, children crying, sunset call to prayer, and motor vehicles. The loudest sounds were produced by insects and mammals, but at frequency bandwidths that did not coincide with the swarm's first harmonic. The sound of mosquito swarms was the only continuous sound in the 100-1000 Hz frequency band (Figure S1).

In addition to these preliminary field recordings at 1 m from the swarm, we found that the first harmonic amplitude of the sound pressure was 10% higher than the background noise (50-Hz smoothed magnitude spectrum), irrespective of which side of the swarm was recorded, i.e. from ground level to the top of a ~3 m-high swarm, and horizontally, on two opposing sides of the swarm at ~3 m from the centre of the swarm. This indicates that the signal-to-noise ratio of the swarm sound can potentially be loud enough to be heard by females at least ~3 m away from the centre of the swarm.

##### Number of males in swarms

In order to infer the sound level of swarms that have more males than those we established under laboratory conditions, we needed to know the range of number of males in natural field swarms. Few studies have investigated the range in numbers of males in mosquito swarms; in Benin, *An. coluzzii* male swarms were typically composed of tens to thousands of males, with a median of ~ 300 males *(S7)*, and in the area of our field study, single sweep-net samples of *An. coluzzii* swarms caught a median of 200 males and a quarter of the samples contained 500–2,500 males *(S8)*, indicating the likelihood that there may be far more males in a swarm than these estimates. As many as 10,000 males in a swarm have been observed in the area of our field study (pers. com. Diabaté). We observed that larger swarms (numerically and spatially) occur in areas and times of year when *An. gambiae s.l.* population densities are highest, especially in the peak of irrigated rice growing during the wet season, and in non-irrigated areas during drier periods, swarms are regularly composed of 20-30 individuals at their peak *(S9)*.

The 70-male swarm used for the laboratory assay is, therefore, realistic, but relatively small compared to the variation observed in the field, and the hearing range prediction based on a 300-male swarm may be considered a typical case.

**Swarm radius as a function of number of males**

A swarm composed of more mosquitoes will produce a higher sound level, and so the distance at which it is audible will increase, accordingly. However, this relationship only has a meaningful real-world impact on swarm localization if the audible distance increases faster than the swarm radius. If the radius increases faster than the distance at which the aggregation is detectable, a female is likely to hear an individual male swarming at the edge of the swarm sooner or more loudly than the swarm as a whole, because particle velocity increases rapidly at close-range of an individual mosquito. For this reason, information on how a swarm radius changes with the number of males is important for the interpretation of our results.

Several studies have investigated qualities of mosquito swarms e.g., the relationship between the marker size and swarm dimension *(S10, S11, S12)*, between the number of males and the marker size *(S11)* or the marker type *(S13)*. In one of our previous studies, the relationship between the number of males and the swarm dimension, given a visual marker, was quantitatively measured *(S9)*. *Anopheles gambiae s.l.* swarms composed of 10 to 50 males in Mali were observed to conform to a bell-shaped distribution of male density over the swarm centre, with a rapid decrease in the number of individuals with distance to the swarm centroid (20% of the swarm's individuals were within a radius of 20 cm of the centre, ~70-90% within 40 cm, 98% within 1 m). Thus, the first effect of increasing the number of males in the swarm is to increase male density in the swarm centre and not throughout the entire volume of the swarm.

Figure S2 uses the data of five swarms of *An. coluzzii* and seven swarms of *An. gambiae s.s.* from *(S9)* to predict swarm radius as a function of the total number of males and of two 'layers' of a swarm (50% and 95% of the most central males), with a random intercept and slope model to predict the radius of swarms consisting of greater number of males, up to the order of thousands. We consider the swarm radius to be defined by the radius of the sphere which encompasses 95% of the males nearest the swarm centroid. The results for *An. coluzzii* are consistent with observations of swarms with thousands of males which are usually < 1 m in radius *(S13)*. For *An. coluzzii*, the predicted mean swarm radius is 0.5±0.1 m for 95% of 1,000 swarming males (0.20±0.05 m for 50% of them) and 0.6±0.1 m for 95% of 10,000 males (0.21±0.05 m for 50% of them), representing a steep increase in density of swarming males, especially in the swarm centre (Figure S2).The swarm radius of an *An. gambiae s.s.* swarms is slightly larger for small swarms, but the predicted radius for large swarms is much larger (Figure S2). The prediction has to be taken with caution for the greatest number of males.

#### Supplementary figures

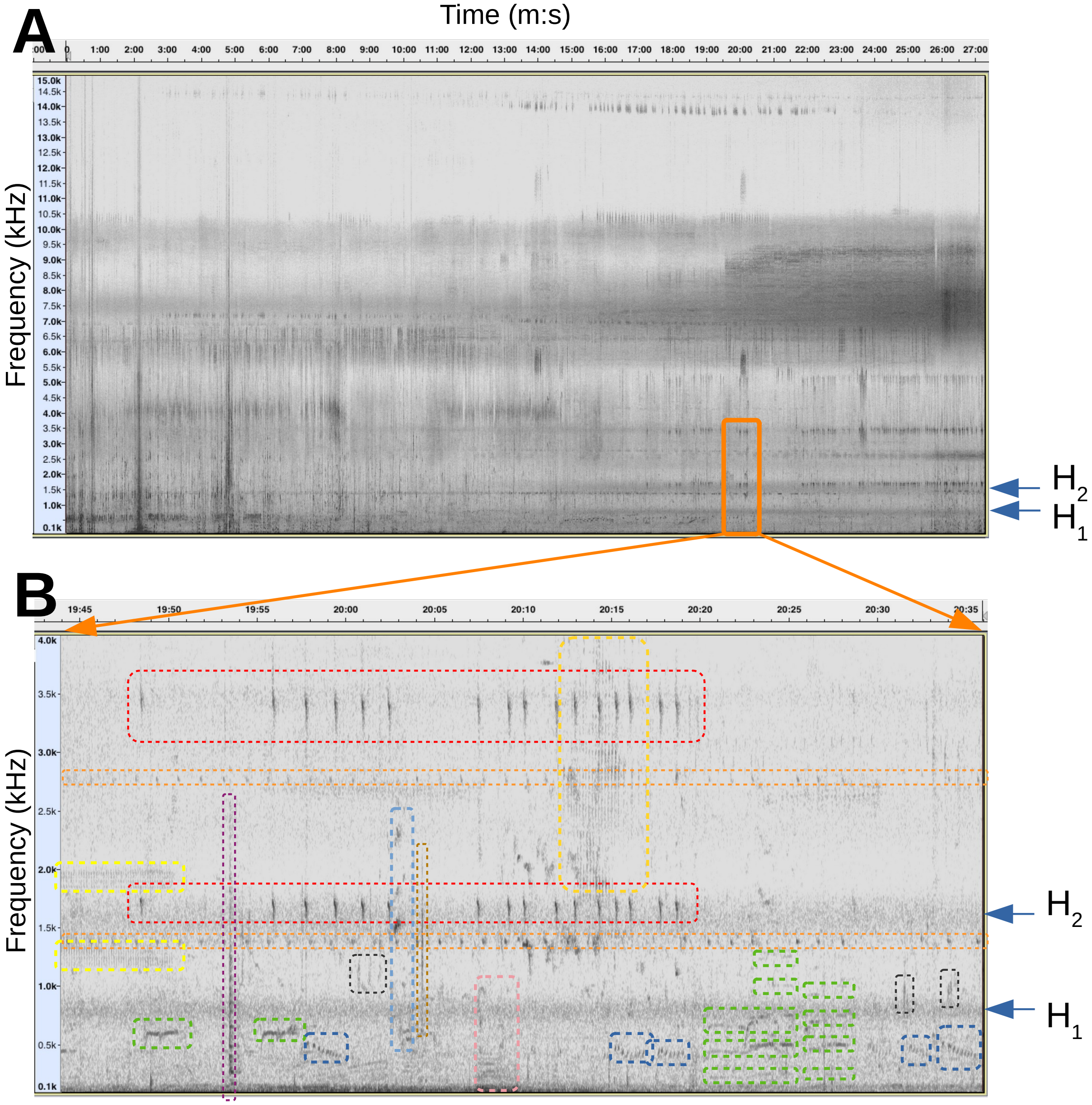


**Figure S1. Spectrogram analysis of a typical field soundscape.** Spectrogram analysis of background sounds in the field over 0-4 kHz in Bama VK5 village, 5 October 2017. X axis represents time, Y axis is the sound frequency and greyscale corresponds to the sound level at a given time and frequency.

**(A)** The spectrogram shows evening sound activity 18h05 - 18h33 (X axis), between 0.1 and 15 kHz (Y axis). Mosquitoes started to swarm ~ 1 min before sound recording began and they start being visible on the spectrogram (i.e. emerging from background sound noise) in the second half of the recording (first and second harmonic frequencies are shown by arrows, H1 and H2, respectively). No sound activity was visible between 15 kHz and 20 kHz, so this frequency range was not displayed. The orange rectangle shows the selected time-frequency area for (B).

**(B)** 52-s spectrogram zoomed-in from orange selection shown in (A). The two arrows show the first and second harmonics of the swarming mosquitoes' wingbeat frequencies. Each colored rectangle shows a different non-mosquito sound source. At one metre from the swarm, the mosquitoes' first harmonics were the only sound-source in this frequency band, while the second harmonics were mixed with a range of bat and frog calls (red, orange, clear yellow, brown) or child voice (light blue). Gold yellow and dark blue correspond to bird sounds; green: mosque call to prayer; pink: motor; purple: cooking metal dish; black: frogs.  Mosquito sounds appear to be the only steady sound in the 100-1000Hz frequency band. Third and fourth harmonic levels are almost invisible.

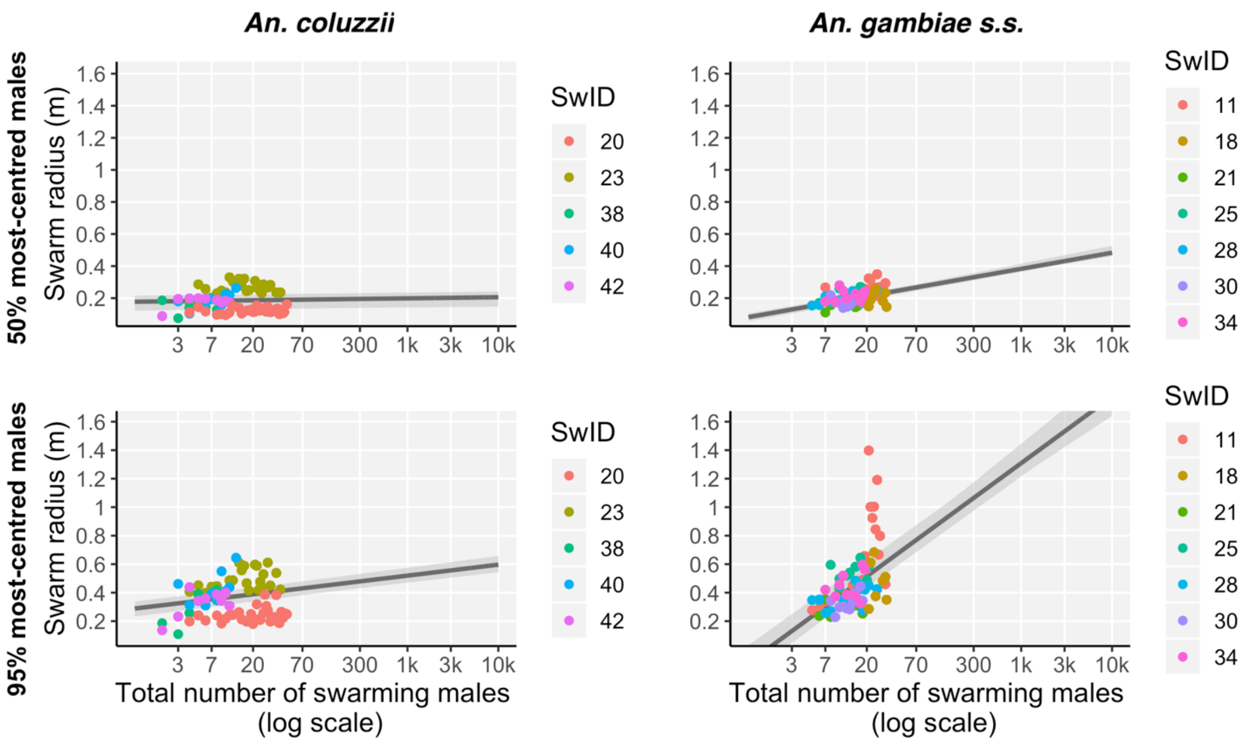


**Figure S2. Radius of wild swarms of *An. coluzzii* and *An. gambiae s.s.* in Mali**. Distance between swarm's centroid and the 50% (top row) or 95% (bottom row) closest males (data from *(S9)*). For each column, color corresponds to a different swarm (left column: *An. coluzzii*.; right column: *An. gambiae s.s.*). Each point of each color corresponds to a measurement made at a different time. The thick black line and grey panel are the regression line of all swarms (random intercept model, *lmer* R function) and the associated 95%-CI. Related to Figure 5.

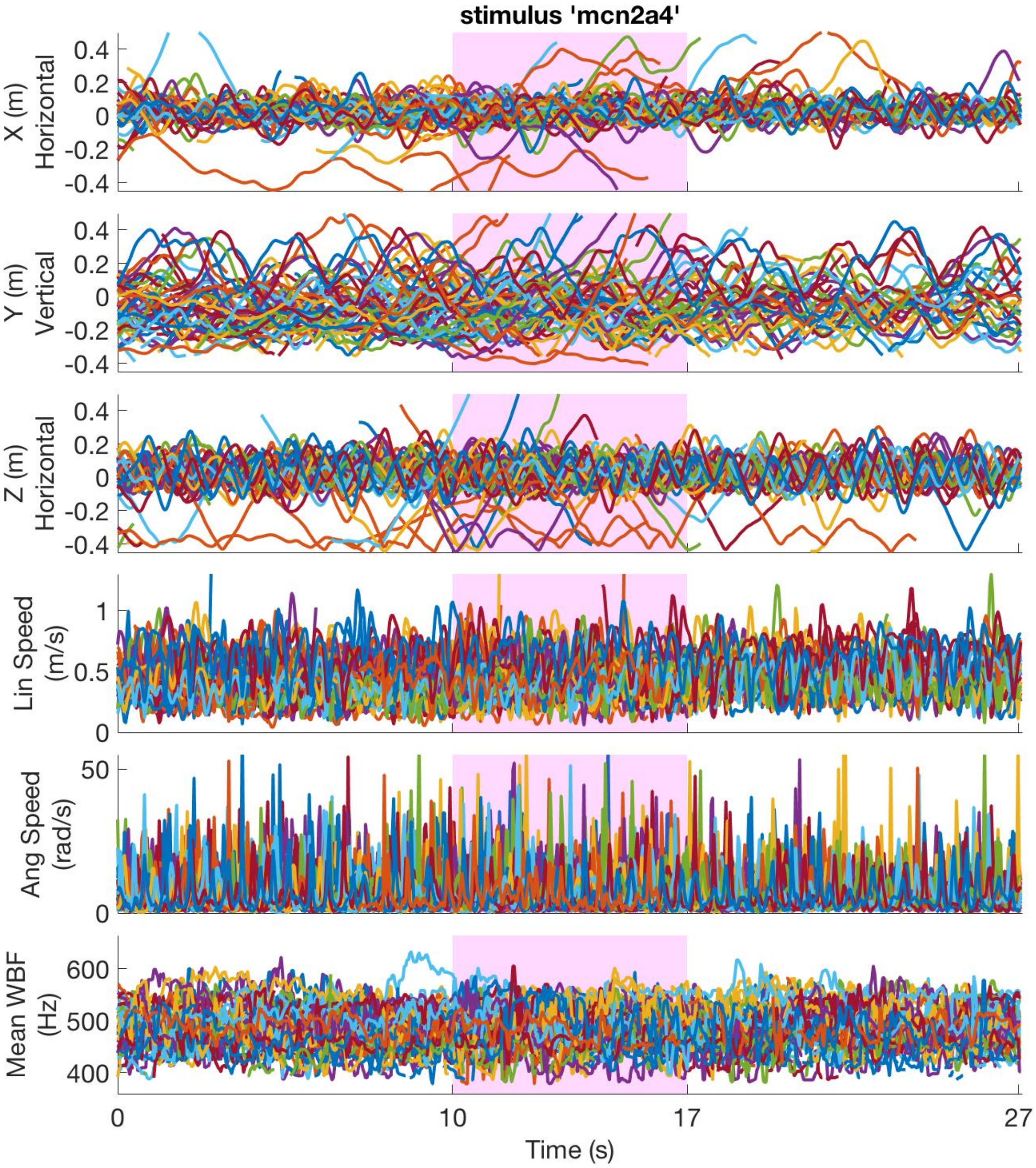


**Figure S3. Superimposed individual flight and sound responses of females to sound stimuli.** *Anopheles coluzzii* response to highest sound-level sound-stimulus over 27 s of recording. Stimulus was played-back 10 s from beginning of flight recording and lasted 7 s (red rectangular shading). First five rows show flight parameters (relative 'XYZ' position, plus linear and angular flight speeds). 'Z' dimension represents relative distance to the speaker (located 0.9 m from *Z*=0). Last row shows mean wingbeat frequency (WBF) of 1st harmonic. Each coloured line represents a different mosquito, and all mosquitoes are plotted for this particular sound stimulus.

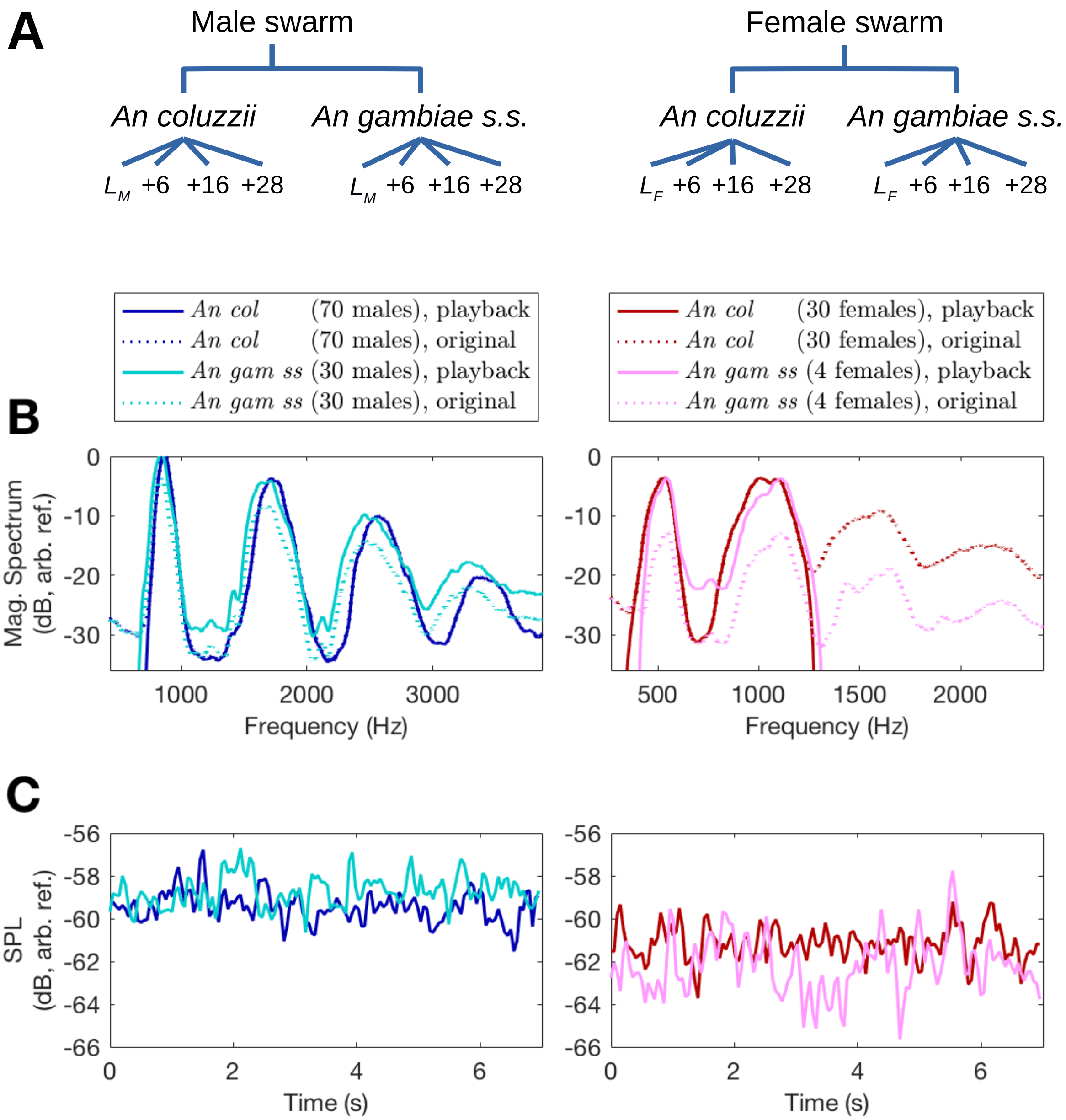


**Figure S4. Spectral and temporal properties of sound stimuli for both sexes and both species.** Spectral and temporal properties of sound stimuli of male *An. coluzzii* swarm (Sound S1), male *An. gambiae s.s.* swarm (Sound S2), female *An. coluzzii* swarm (sound S3) and female *An. gambiae s.s.* swarm (Sound S4).

**(A)** Sound stimulus variables: mosquito sex; species; and amount of sound level increase compared to a sex-specific reference level ($L_M$ or $L_F$).

**(B)** Magnitude spectrum, computed over 7 s and averaged over 50-Hz windows, of stimuli of each species (dark colors for *An. coluzzii* and light colors for *An. gambiae s.s.*) and each sex (blue for males and red for females). Dotted lines represent recorded sounds, while solid lines indicate played-back sounds, the level for which was adjusted to one of *An. coluzzii*'s first-harmonics (see Supplementary Methods section 'Generation of sound stimuli'). High-frequencies of the female-swarm sound were filtered out to allow monitoring of the sound harmonics of the males, which otherwise would overlap with the females' higher harmonics (see Supplementary Methods section "Wingbeat parameter extraction from flight sound"). The number of mosquitoes in the original sound stimuli is indicated in brackets in the legend.

**(C)** Root-Mean-Square pressure level (0.1 s time windows with 0.05 s overlap), over the 7-s duration of the stimuli, for each species (dark colors: *An. coluzzii*, light colors: *An. gambiae s.s.*) and each sex (blue: males and red: females).

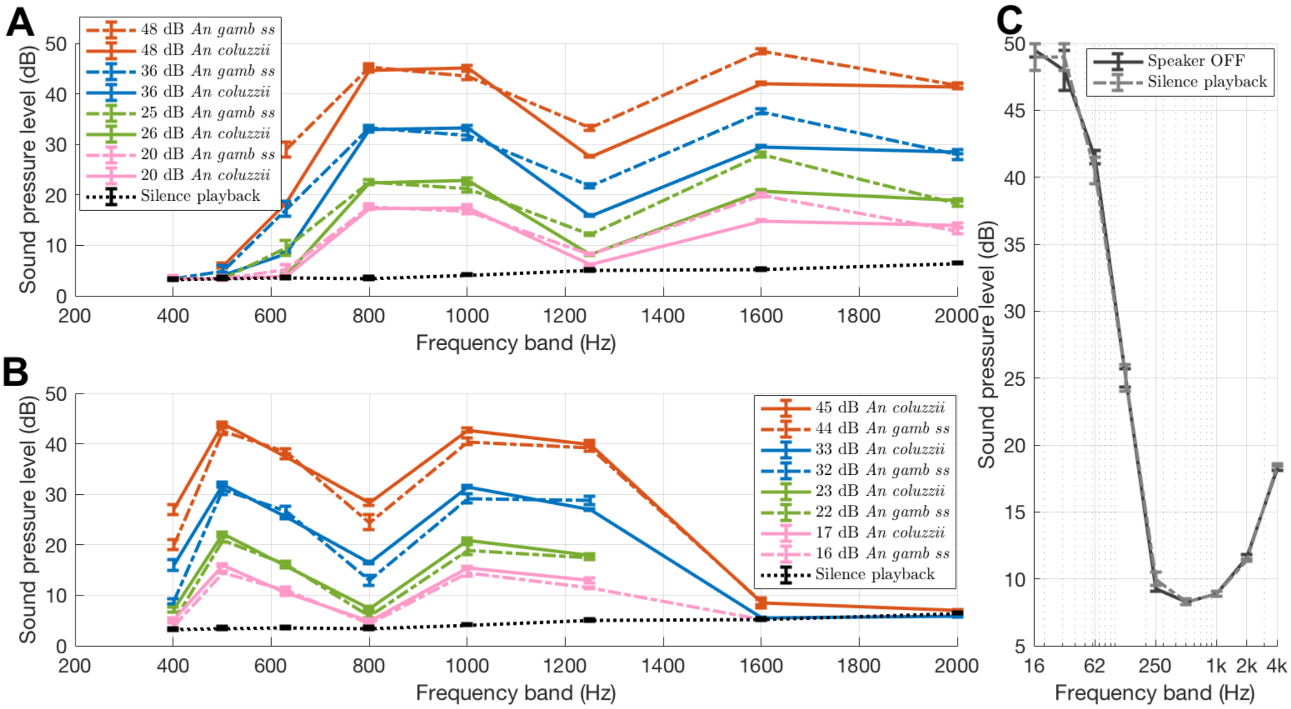


**Figure S5. Calibrated sound-level measurements of played-back swarm sounds and background noise.** SPLs (ref 20 µPa) were measured at the *An. coluzzii* swarming position 1) to estimate the sound level received by the tested mosquito and 2) to know the background noise level of the soundproof chamber. The X-axis values represent the central frequency $f_c$ of the octave or 1/3-octave band filters. Each filter has a lower limit of $2^{-1/6}f_c$ and an upper limit of $2^{1/6}f_c$. For example, X=800 Hz represents the sound pressure level between 713 Hz and 898 Hz. The Y-axis error-bar represents maximum and minimum values measured during stimulus duration under a time constant of 1 s (slow mode, according to IEC 61672-1: 2002).

**(A)** SPL measurements of male-swarm sound stimuli as a function of frequency, limited to a frequency range audible to *An. coluzzii* females *(S2)*, and plotted at 1/3 octave steps reveals first and second harmonics of the swarming-male wingbeat sound. Black dashed line shows sound level when playing-back 'silent' (i.e. just speaker noise), with the same settings as during the experiment. The four solid lines correspond to the sound levels at the mean female's 'XYZ' position during play-back of each of the four *An. coluzzii* male-swarm stimuli (related to four distances from the male-swarm sound image, see Figure 3). Dashed lines represent the same, but for *An. gambiae* sound stimuli.

**(B)** As for A, except related to female sound stimuli. Not all points were measured above 1250 Hz which is of less interest because we filtered out sound frequencies above the second harmonics.

**(C)** SPL measurements (ref 20 µPa) in the soundproof chamber without playback and with 'silence' playback, along octave bands from 16 Hz to 4 kHz. The black dashed line shows the sound level when playing-back silence (i.e. speaker noise), while the plain black line corresponds to when the speaker is off (i.e. showing noise floor of sound-proof room). Both are the same, i.e. they show that the speaker has a very low noise level enabling us to playback low level sounds.

#### Supplementary tables

| Index | SPL at oscillatory distances from female(s) to speaker due to swarming behaviour (room mode effect included) | Estimated distances between the female(s) and the sound-source image of a male swarm | | | | |
|---|---|---|---|---|---|---|
| | | $r_i$ (calculated from $L_i$) | $r_{i,j}$ (calculated from $r_i$) j number of swarming males | | | |
| i | SPL (measured) $L_i$ (dB) | 70-male swarm $r_i$ | 300-male swarm $r_{i,300}$ | 1500-male swarm $r_{i,1500}$ | 6,000-male swarm $r_{i,6000}$ | 10,000-male swarm $r_{i,10k}$ |
| ref | 20±3 | 0.9±0.3 m (recorded case) | 1.9±0.7 m | 4.3±1.5 m | 9±3 m | 11±4 m |
| 1 | 26±3 | 0.5±0.2 m | 1.1±0.4 m | 2.4±0.9 m | 4.9±1.8 m | 6±2 m |
| 2 | 36±3 | 16±6 cm * (37 dB SVL) | 0.3±0.1 m | 0.7±0.3 m | 1.4±0.5 m | 1.9±0.7 m |
| 3 | 48±3 | 4±1 cm * (54 dB SVL) | 8±3 cm * (51 dB SVL) | 19±6 cm * (49 dB SVL) | 0.4±0.1 m | 0.5±0.2 m |

**Table S1. Schematic relationship between 'sound level' and 'distance' for *An. coluzzii* sound stimuli.** Table shows estimated distances from the female(s) to the sound-source image of male-swarm sound, played-back 0.9 m from the centre of the females' swarming area. SPLs were computed from the calibrated SPL measurements (ref 20 µPa) of the two nearest third-octave bands to the wingbeat frequency's first-harmonic. SPL errors were computed by taking into account the oscillating distance between the female(s) and the speaker due to their swarming behaviour above the visual marker (see Figure 2 and Supplementary Methods section 'Sound pressure level'). The distances $r_i$ from the female to the sound-source image of the 70-male swarm sound-stimuli were computed from an acoustic-propagation formulae using $L_i$ and $r_{ref}$ (see Methods equation 4) and the errors were directly derived from SPL errors. The equivalent distance $r_{i,j}$ for a j-male swarm, to result in the same sound pressure level $L_i$, was extrapolated from $r_i$ using another acoustic formula (see Methods equation 5). The asterisk (*) means that the distance should be greater than indicated or the sound particle-velocity level (SVL) should be greater than the SPL as indicated (see Supplementary Methods section 'Relationship between particle-velocity and pressure levels). The SPL measurements of *An. gambiae s.s.* sound stimuli are reported in Table S2; they were close to values for *An. coluzzii,* resulting in similar estimated distances between the female(s) and the sound-source image of a male swarm.

| Subjects exposed to sound stimuli | Sound stimuli | | | | | |
|---|---|---|---|---|---|---|
| **Species and sex** | **Sex** | **Number of swarming individuals** | **Species** | **Number of harmonics** | **Sound level** | |
| | | | | | **Played-back gain of 50Hz-smooth 1st-harmonics** | **SPL measurement of the two 1/3-octave bands closest to the 1st harmonic (dB SPL) at fixed distances from the speaker (0.9 m)** |
| NA | Silence playback | | | | NA | 6.9±0.3 |
| *An. coluzzii* female | Male | Group (~70) | *An. coluzzii* | all | $L_M$, related to natural SPL 90cm away from the speaker | 20±3 |
| | | | | | $L_M$+6dB | 26±3 |
| | | | | | $L_M$+16dB | 36±3 |
| | | | | | $L_M$+28dB | 48±3 |
| | | Group (~30) | *An. gambiae* | | $L_M$ | 20±3 |
| | | | | | $L_M$+6dB | 25±3 |
| | | | | | $L_M$+16dB | 36±3 |
| | | | | | $L_M$+28dB | 48±3 |
| *An. coluzzii* male | Female | Group (~30) | *An. coluzzii* | 2 | $L_F$, related to natural SPL 90cm away from the speaker | 17±3 |
| | | | | | $L_F$+6dB | 23±3 |
| | | | | | $L_F$+16dB | 33±3 |
| | | | | | $L_F$+28dB | 45±3 |
| | | Group (~4) | *An. gambiae* | | $L_F$ | 16±3 |
| | | | | | $L_F$+6dB | 22±3 |
| | | | | | $L_F$+16dB | 32±3 |
| | | | | | $L_F$+28dB | 44±3 |

**Table S2. Description of stimulus loudness at fixed distances from the speaker**. This table gives the sound level of all played-back sound stimuli at fixed distances to the speaker, according to two different approaches. The first is the relative signal gain added to the played-back sound, ranging from +0 dB to +28 dB, with 0 dB relative to the natural sound level 0.9 m away from either a 70-male *An. coluzzii* swarm ($L_M$) or a 30-female *An. coluzzii* swarm ($L_F$) (see Supplementary Methods section 'Generation of sound stimuli' for the calculation of $L_M$ and $L_F$). The gain of played-back sound of the *An. gambiae* swarm sound stimuli were corrected to be the same as that of the *An. coluzzii swarm,* to balance the different number of mosquitoes in the swarms of each species. The second approach to describe the sound level is to measure a calibrated sound pressure level (SPL ref 20 µPa) of the played-back sound stimulus at the mosquito's mean location in the frequency range of the opposite-sex's harmonics audible by the mosquito (see Supplementary

Methods section ‘Wingbeat parameter extraction from flight sound’). SPL errors were estimated using minimum and maximum sound pressure levels over time.

| | | Explanatory variables | | | | | |
|---|---|---|---|---|---|---|---|
| Response variable (diff) | | SPL:species | | species | | SPL | |
| | Median computation duration on either side of the stimulus onset (s) | $\chi_1^2$ | p | $\chi_1^2$ | p | $\chi_1^2$ | p |
| Wingbeat frequency | 1 | 0.29 | 0.59 | 1.1 | 0.29 | 1.1 | 0.28 |
| | 7 | 0.07 | 0.79 | 0.023 | 0.88 | 3.2 | 0.074 |
| Height | 1 | 5.9 | 0.015* | 8.7 | 0.0032** | 3.1 | 0.076 |
| | 7 | 0.60 | 0.44 | 0.47 | 0.49 | 0 | 1 |
| Distance to speaker | 1 | 1.3 | 0.25 | 2.2 | 0.14 | 4.6 | 0.032* |
| | 7 | 0.18 | 0.67 | 0.29 | 0.59 | 0.15 | 0.70 |
| Angular speed | 1 | 0.59 | 0.44 | 3.8 | 0.051 | 3.7 | 0.055 |
| | 7 | 0.51 | 0.47 | 1.0 | 0.31 | 0.010 | 0.92 |
| Linear speed | 1 | 0.31 | 0.58 | 0.0004 | 0.98 | 0.13 | 0.72 |
| | 7 | 1.9 | 0.16 | 0.25 | 0.62 | 1.6 | 0.21 |

**Table S3. Statistical results from additional parameters computed from female wingbeat frequency or flight trajectory.** Response parameters were calculated both during the sound stimulus playback and just before the playback for the same duration (1 or 7 s), and then were differentiated. LRT $\chi^2$ was calculated between selected model pairs for which stepwise removal of terms was used. The distance-to-speaker parameter showed a significant effect of SPL but the associated distributions were not significantly different from the intercept at the $p$=0.05 level, including the loudest sound stimulus, 48 dB distribution (one-sample $t(22)$=-1.51, BH-corrected $p$=0.29, mean=-0.02 rad/s). The effects of SPL:species over the height parameter was also considered as false positive because only one of the smallest SPL distribution was associated to a significant difference from the intercept at the $p$=0.05 level (*An. coluzzii* 26 dB stimulus: one-sample $t(9)$=4.16, $p$=0.020, mean=0.04 m; *An. gambiae* 20 dB stimulus: $t(11)$=3.13, $p$=0.038, mean=0.05 m). Since the effect of SPL:species was considered as false positive and since there was no SPL effect, the species effect was consequently considered as false positive. Moreover, if considering the large number of tested parameters (additional BH-correction for the 10 parameters), no distributions were significantly different from the intercept.

#### Other supplementary files

**Sound S1 (Sound-S1.mp3)**. Sound stimulus recording of the 70-male *An. coluzzii* (7 s) before any filtering and level adjustment. Related to Figure S4B (dotted clear blue line).

**Sound S2 (Sound-S2.mp3)**. Sound stimulus recording of the 30-male *An. gambiae s.s.* (7 s) before any filtering and level adjustment. Related to Figure S4B (dotted dark blue line).

**Sound S3 (Sound-S3.mp3)**. Sound stimulus recording of the 30-female *An. coluzzii* (7 s) before any filtering and level adjustment. Related to Figure S4B (dotted clear red line).

**Sound S4 (Sound-S4.mp3)**. Sound stimulus recording of the 4-female *An. gambiae s.s.* (7 s) before any filtering and level adjustment. Related to Figure S4B (dotted dark red line).

**Video S1 (Video-S1.mp4)**. Audio-video recording of the *An. coluzzii* female exposed to the loudest *An. coluzzii* male sound (10-s silence + 7-s sound exposition + 10-s silence). Related to Figure 2.

**Video S2 (Video-S2.mp4).** Audio-video recording of the *An. coluzzii* males exposed to the loudest *An. gambiae s.s.* female sound (10-s silence + 7-s sound exposition + 10-s silence). Related to Figure 2.
